# BayFlux: A *Bay*esian method to quantify metabolic *Flux*es and their uncertainty at the genome scale

**DOI:** 10.1101/2023.04.19.537435

**Authors:** Tyler W. H. Backman, Christina Schenk, Tijana Radivojevic, David Ando, Janavi Singh, Jeffrey J. Czajka, Zak Costello, Jay D. Keasling, Yinjie Tang, Elena Akhmatskaya, Hector Garcia Martin

## Abstract

Metabolic fluxes, the number of metabolites traversing each biochemical reaction in a cell per unit time, are crucial for assessing and understanding cell function. ^13^C Metabolic Flux Analysis (^13^C MFA) is considered to be the gold standard for measuring metabolic fluxes. ^13^C MFA typically works by leveraging extracellular exchange fluxes as well as data from ^13^C labeling experiments to calculate the flux profile which best fit the data for a small, central carbon, metabolic model. However, the nonlinear nature of the ^13^C MFA fitting procedure means that several flux profiles fit the experimental data within the experimental error, and traditional optimization methods offer only a partial or skewed picture, especially in “non-gaussian” situations where multiple very distinct flux regions fit the data equally well. Here, we present a method for flux space sampling through Bayesian inference (BayFlux), that identifies the full distribution of fluxes compatible with experimental data for a comprehensive genome-scale model. This Bayesian approach allows us to accurately quantify uncertainty in calculated fluxes. We also find that, surprisingly, the genome-scale model of metabolism produces narrower flux distributions (reduced uncertainty) than the small core metabolic models traditionally used in ^13^C MFA. The different results for some reactions when using genome-scale models vs core metabolic models advise caution in assuming strong inferences from ^13^C MFA since the results may depend significantly on the completeness of the model used. Based on BayFlux, we developed and evaluated novel methods (P-^13^C MOMA and ROOM) to predict the biological results of a gene knockout, that improve on the traditional MOMA and ROOM methods. We provide an open source Python implementation of BayFlux at https://github.com/JBEI/bayflux.

**Author summary:** ^13^C MFA practitioners know that modeling results can be sensitive to minor modifications of the metabolic model. Certain parts of the metabolic model that are not well mapped to a molecular mechanism (*e.g.* drains to biomass or ATP maintenance) can have an inordinate impact on the final fluxes. The only way to ascertain the validity of the model is by checking that the result does not significantly differ from previously observed flux profiles. However, that approach diminishes the possibility of discovering truly novel flux profiles. Because of this strong dependence on metabolic model details, it would be very useful to have a systematic and repeatable way to produce these metabolic models. And indeed there is one: genome-scale metabolic models can be systematically obtained from genomic sequences, and represent all the known genomically encoded metabolic information. However, these models are much larger than the traditionally used central carbon metabolism models. Hence, the number of degrees of freedom of the model (fluxes) significantly exceeds the number of measurements (metabolite labeling profiles and exchange fluxes). As a result, one expects many flux profiles compatible with the experimental data. The best way to represent these is by identifying all fluxes compatible with the experimental data. Our novel method BayFlux, based on Bayesian inference and Markov Chain Monte Carlo sampling, provides this capability. Interestingly, this approach leads to the observation that traditional optimization approaches can significantly overestimate flux uncertainty, and that genome-scale models of metabolism produce narrower flux distributions than the small core metabolic models that are traditionally used in ^13^C MFA. Furthermore, we show that the extra information provided by this approach allows us to improve knockout predictions, compared to traditional methods. Although the method scales well with more reactions, improvements will be needed to tackle the large metabolic models found in microbiomes and human metabolism.

## Introduction

Synthetic biology enables us to bioengineer cells for synthesis of novel valuable molecules such as renewable biofuels or medical drugs [1–5], but its full potential is hindered by our inability to predict biological behavior [6, 7]. We can engineer DNA changes faster than ever, and we can measure the impact of these genetic changes in more detail than ever through an increasing amount of functional genomics data. But the availability of all these advances does not necessarily translate into better predictive capabilities for biological systems: converting the collected data into actionable insights to achieve a given goal (*e.g.*, higher bioproduct yields) is far from trivial or routine.

Metabolic fluxes (*i.e.*, the number of metabolites traversing each biochemical reaction per unit time per a given amount of biomass) are crucial to predict and understand biological systems because they map how carbon and electrons flow through metabolism to enable cell function. Flux analysis, for example, has been used to improve biofuel production [8], contextualize multiomics data [9], and provide insights into multi-species relationships [10].

Although there are several popular methods for studying metabolic fluxes [11–13] ^13^C MFA is considered to be the gold standard to measure metabolic fluxes. Metabolic Flux Analysis (MFA [11], as opposed to ^13^C MFA) works by using measurements of the exchange fluxes (fluxes coming in and out of the cell) to fully constrain a small core metabolic network (with less degrees of freedom than measurements). Flux Balance Analysis (FBA [12]) uses a comprehensive metabolic network (a genome-scale model, or GSM) that encompasses all reactions encoded in the genome, constrains it using the exchange fluxes, and then finds the fluxes corresponding to the highest growth rates. Traditional ^13^C MFA [13] works by using exchange fluxes as well as data from ^13^C labeling experiments to find which flux profile (*i.e.*, set of fluxes for every reaction in the model) best describes the data for a small metabolic model. Genome-scale ^13^C MFA uses the same data types but finds the flux profile for a (much larger) genome-scale metabolic model, often containing thousands of reactions [14, 15]. FBA can produce good results when the cells are under selection for maximum growth [16] but is less useful when that is not the case (*e.g.* human cells or engineered bacterial strains). ^13^C MFA is considered the most accurate approach to measure metabolic fluxes but relies on expensive and time consuming ^13^C experiments which are nontrivial to do in a high-throughput fashion. Both methods have been used successfully to guide bioengineering processes [17, 18].

The optimization approach used to date in ^13^C MFA to determine fluxes [19–22] shows several limitations, particularly in characterizing the full distribution of fluxes compatible with the data. For example, the results from small core metabolic models can be very sensitive to the modification of apparently innocuous components of the model; if genome-scale models are in use the system shows more degrees of freedom (fluxes) than experimental data and we expect many flux profiles to be compatible with the experimental data. Furthermore, errors in determining ^13^C labeling for a single metabolite can completely derail flux calculation. This optimization approach also often depends on expensive commercial solvers, and is hard to parallelize. Finally and most importantly, uncertainty quantification relies on confidence intervals estimated with help of frequentist statistics. Such intervals depend on a given experimental outcome and may result in misinterpretation of the flux uncertainty, as has been noted by others before us [23]. For example, the solution space may have an area of poor fit to the data between two distinct regions of excellent fit (*e.g.* non-gaussian fitness distribution), such that a single point does not meaningfully represent the experimental data.

Here, we present a method (BayFlux) that rigorously identifies the full distribution of flux profiles compatible with experimental data for a genome-scale model (Fig. 1). The result provided to the end user is a “probability distribution” of possible fluxes, which faithfully reports the full uncertainty due to experimental error, and any potential model data incompatibilities. To achieve this, we combine *both* Bayesian inference and Markov Chain Monte Carlo (MCMC) methods to sample flux space, as informed by ^13^C labeling and flux exchange data. Just Monte Carlo sampling by itself provides all fluxes compatible with experimental data instead of just the fluxes that best fit the available data [24–27]. However, while this approach offers much needed flux uncertainty quantification [28], it fails when the data used is inconsistent (*e.g.* fluxes coming in and out of the cell are not mass balanced). Bayesian inference can address this complication because it is is based on a probabilistic interpretation that ensures a systematic approach to manage data inconsistencies and update flux probability distributions as more data becomes available [29–32]. Combining Monte Carlo flux sampling with Bayesian statistics hence provides reliable flux uncertainty quantification in a way that scales efficiently as more data become available [23]. Unlike previous approaches [23, 30, 33], our approach enables flux uncertainty quantification for genome-scale models rather than only for small central carbon metabolism models [23], and helps trace the origin of flux uncertainty directly to the physical measurements of metabolite labeling. Using genome-scale models for modeling metabolism provides a comprehensive understanding of all metabolic fluxes in a cell [14, 15], as well as standardizing the application of ^13^C MFA. Furthermore, MCMC Bayesian inference is easier to parallelize, use with large datasets, and integrate with heterogenous data sources (*e.g.* multiomics data [31]).

**Fig 1.**
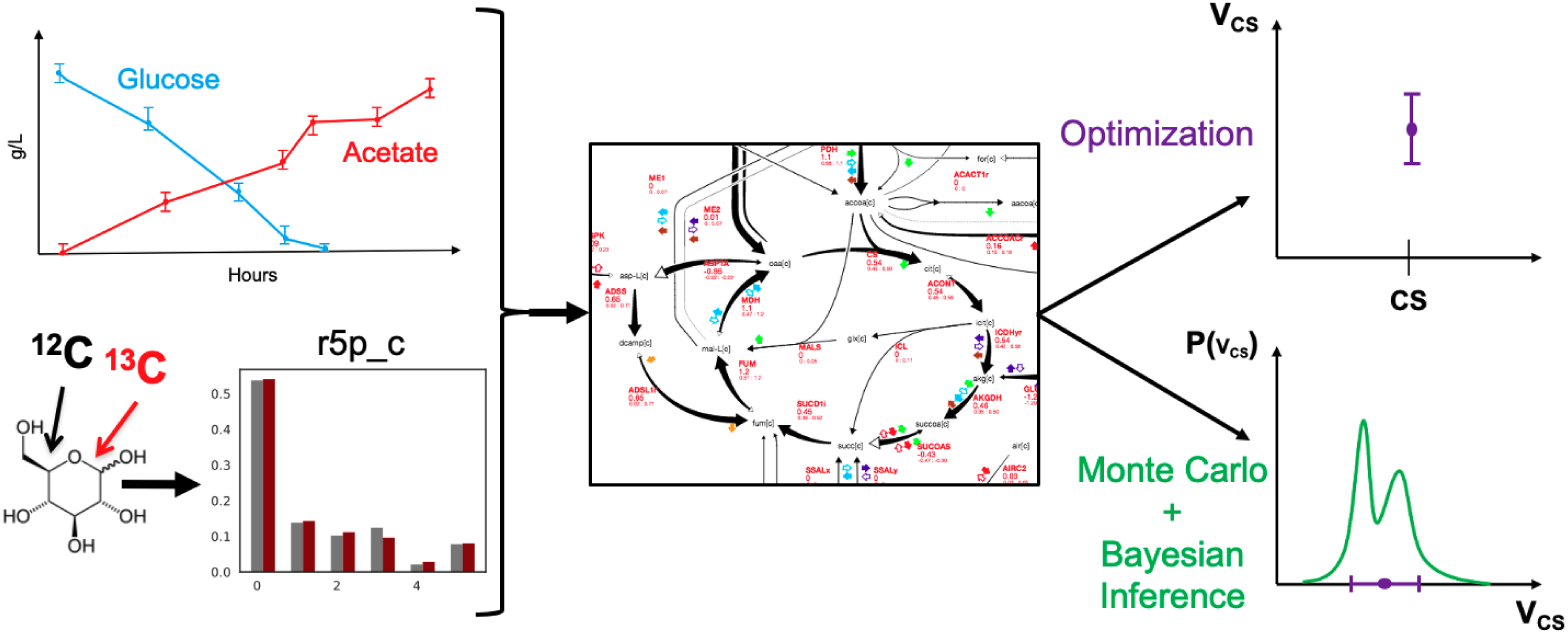
A new approach to calculating metabolic fluxes. The state of the art in measuring metabolic fluxes (^13^C MFA) involves using ^13^C experimental data and extracellular metabolite concentration data to find the fluxes that best fit the data (optimization approach) for a core metabolic model. However, core metabolite models only represent a small fraction of all possible reactions and involve simplifications that can have an inordinate impact on the calculated fluxes. Genome-scale models can be systematically derived from the organism genome and represent a comprehensive description of metabolism, but also display more degrees of freedom (reactions) than measurements. This mismatch results in several flux profiles being compatible with the experimental data, which are badly represented by a single flux solution, even if coupled with a confidence interval. BayFlux uses Bayesian Inference and Monte Carlo sampling to provide the full distribution of fluxes compatible with the experimental data.

We showcase BayFlux by performing the first flux sampling for a genome-scale model as constrained by ^13^C data, and demonstrating more informative flux profiles than can be achieved with non-Bayesian, optimization-based approaches. Using an *E. coli* model and data set [34], we show that BayFlux produces results that are compatible with optimization results, and also offer valuable uncertainty quantification information. This uncertainty quantification allows us to show that optimization approaches can overestimate flux uncertainty by representing it through only two numbers: the upper and lower confidence intervals. We find that genome-scale models of metabolism result in narrower flux distributions than the small core metabolic models that are traditionally used in ^13^C MFA. Based on BayFlux, we develop and evaluate novel methods (P-^13^C MOMA and ROOM) to predict the biological results of a gene knockout, that improve on FBA-based MOMA and ROOM methods. Finally, we find that BayFlux scales well as more reactions are added, but efficiency improvements will be needed to sample the very large metabolic models required for microbiomes or human metabolism. We also discuss ideas for future improvement.

## Results and discussion

### BayFlux Monte Carlo sampling results are compatible with optimization results

We obtain compatible results when calculating fluxes through the sampling and optimization approaches for the same core metabolic model (Figs. 2,3). We compared our new BayFlux sampling approach with the optimization approach via a classical ^13^C MFA tool: 13CFLUX2 [34], which leverages the IPOPT (www.coin-or.org/ipopt) and NAG C (www.nag.co.uk) mathematical optimization libraries. For this comparison, we used the *E. coli* data and core metabolic model employed in the demonstration of 13CFLUX2: measurements of glucose uptake, growth rate, and the labeling of eleven central carbon intracellular metabolites for an *E. coli* MG1655 strain grown in glucose-limited continuous culture, and a model comprised of 66 reactions and 37 metabolites describing central carbon metabolism (see Core Metabolic Model 1 below). Flux profiles for the best fit and best sample (*i.e.* highest likelihood) from both methods are very similar (Fig. 2). Notice that the 13CFLUX2 fluxes are always enclosed by the BayFlux probability distributions, and close to the BayFlux best sample (Fig. 3). However, BayFlux provides much more information on flux uncertainty. BayFlux reports the full flux probability distribution, which is rarely uniform over the optimization confidence interval. Furthermore, the corresponding metabolite labeling patterns (mass distribution vectors or MDVs) are virtually identical between 13CFLUX2 and BayFlux (Fig. S1). To enable direct point comparison, we compare the point solution from 13CFLUX2, to the highest posterior probability sample obtained during BayFlux sampling (Fig. 2). These results hold for fifteen instances of flux profiles that were obtained through both methods (Figs S2 and S3).

**Fig 2.**
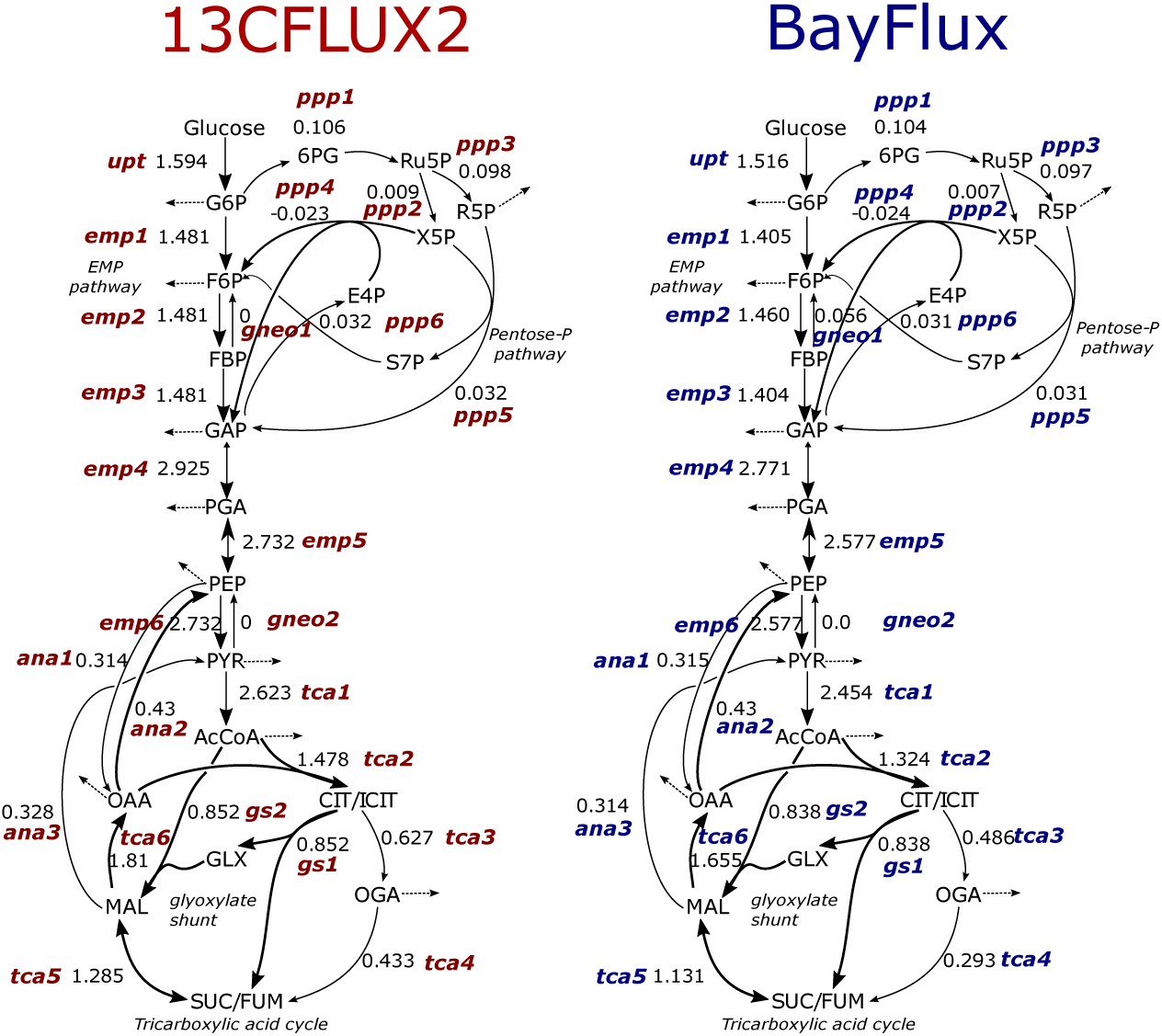
Core flux profiles for simplified metabolic models obtained through BayFlux (sampling, in blue) and 13CFLUX2 (optimization, in red) are similar for the best fit (13CFLUX2) and best sample (BayFlux). The best sample (e.g. highest posterior probability) from BayFlux (for ten million samples) is here compared with the best fit obtained from 13CFLUX2. Results for best fits and samples are similar, but BayFlux offers more information regarding the nuances of the full distribution of fluxes compatible with the experimental data (Fig. 3). All the fluxes are in units of mM/gDW/h. Comparisons for other flux profiles can be found in Fig. S2. Reaction names correspond to Core Metabolic Model 1 (see Materials and Methods).

**Fig 3.**
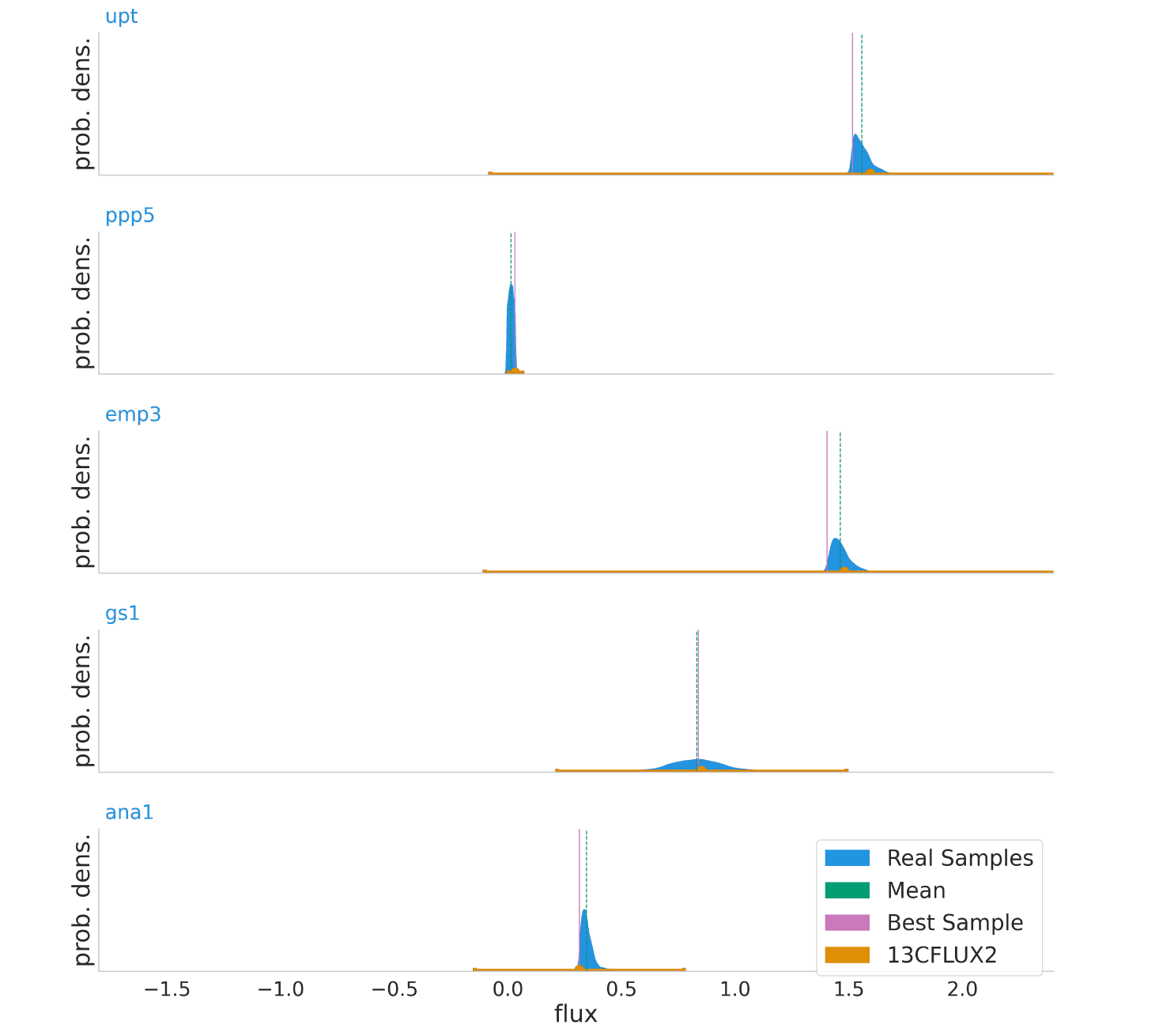
Fluxes obtained from BayFlux using a flux sampling approach are compatible with the optimization results from 13CFLUX2, but offer more information. Whereas the optimization approach only provides the best fit and confidence intervals, BayFlux supplies the probability distribution of all fluxes compatible with the experimental ^13^C data (Fig. 1). Probability densities (blue), best sample (vertical magenta line), and mean (vertical green line) from BayFlux for ten million flux samples are shown vs. 13CFLUX2 best fit with confidence intervals (in orange) for 5 out of 66 fluxes (see Fig. 2 for best fits and best samples for a greater number of reactions). Reaction names correspond to Core Metabolic Model 1 (see Materials and Methods).

The rigorous uncertainty quantification provided by BayFlux shows that the 13CFLUX2 confidence intervals often grossly overestimate flux uncertainty (Fig. 3). For example, fluxes ‘upt’ and ‘emp3’ show a probability density that extends over a range that is an order of magnitude smaller than the corresponding confidence interval. Hence, these fluxes can be determined more accurately through BayFlux than it is suggested by the optimization approach.

### Genome-scale models produce narrower flux distributions than core metabolic models

Flux sampling (via BayFlux) for the genome-scale model of metabolism generally produces narrower distributions of fluxes compatible with the experimental data than the small core metabolic models that are traditionally used in ^13^C MFA (Fig. 4). This means that fluxes are more accurately determined by using genome-scale models than small core models. For this comparison, we used the *E. coli* data and core metabolic model previously published by Toya *et al.* [35] (63 reactions and 47 metabolites describing central carbon metabolism, see Core Metabolic Model 2 below), and as the genome scale model we used the *E. coli* genome-scale model combining iAF1260 [36] and imEco726 [14] models (see Genome-scale *E. coli* atom mapping model below). This finding is surprising because one would expect that the extra degrees of freedom provided by the several thousand reactions in the genome-scale model (as compared to the ∼60 in the core model) would allow for many more distinct flux profiles to meet stoichiometric constraints and labeling data. However, it seems that central metabolic fluxes are well constrained by metabolite labeling, and the addition of several hundreds of extra reactions only add draws of cofactors that further constrain core fluxes. This is consistent with the known bow tie structure of metabolism [37]. Exceptions to this observation involve reactions F6PA (fructose 6-phosphate aldolase), DHAPT (dihydroxyacetone phosphotransferase), and PYK (Pyruvate kinase), which show greater uncertainty (*e.g.* broader probability distrbution peaks) for the genome-scale model than the core model (Fig. 4).

**Fig 4.**
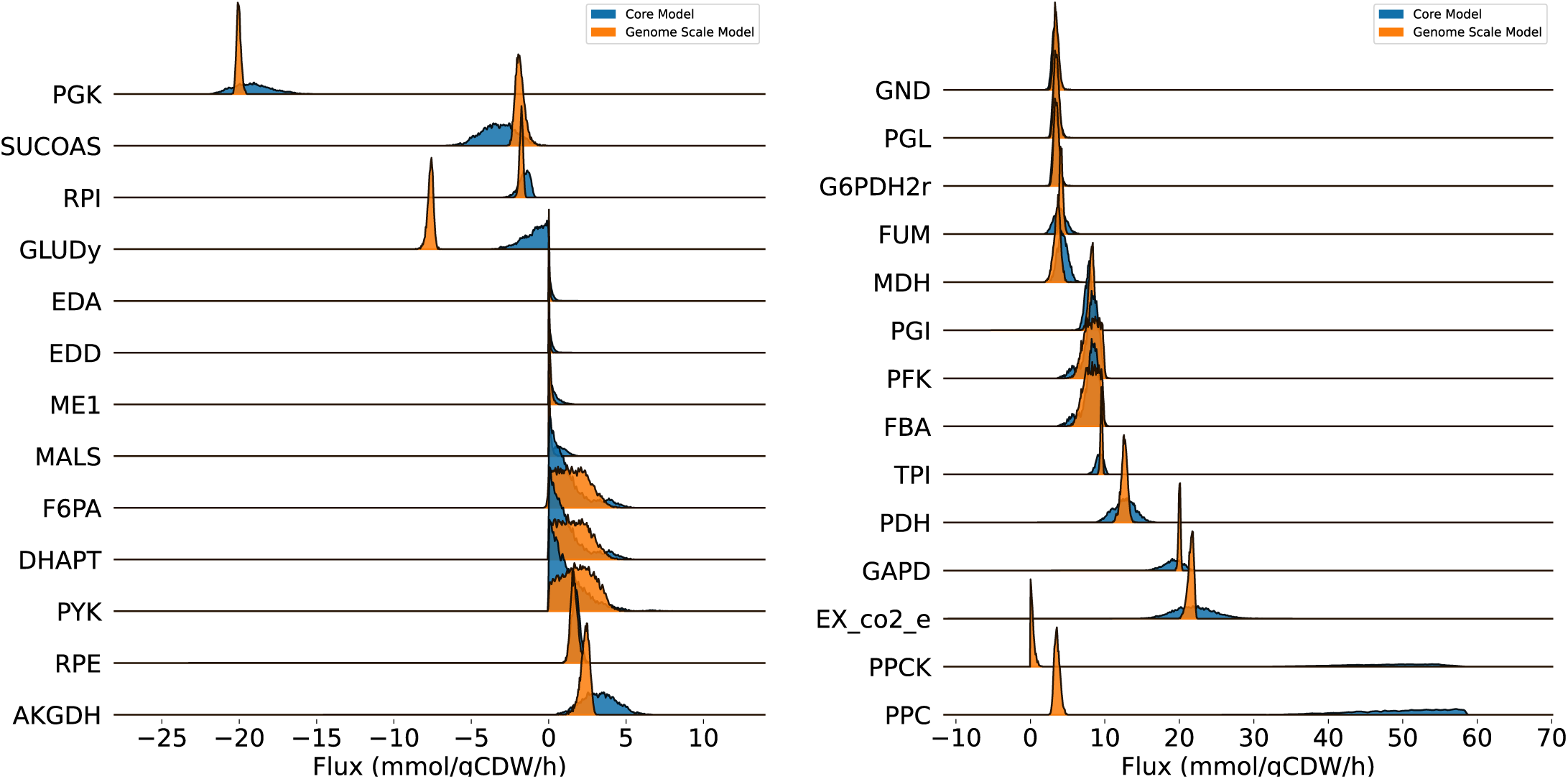
Using genome-scale models produces more biologically meaningful solutions. The results obtained from BayFlux with a core metabolic model (blue) are compared with those obtained from a genome-scale model (orange). Using a genome-scale model produces a narrower flux distribution (higher certainty posterior probability distributions), as informed by a greater amount of biological knowledge encoded in the genome-scale model. Notice too, that certain reactions display very different averages. For example, GLUDY shows very different averages for the genome-scale and core metabolic models, advising caution in assuming strong inferences from ^13^C MFA since the results may depend significantly on the model used. Additionally, several of the probability distributions are non-Gaussian, which can only be meaningfully represented as a full distribution rather than a point or interval. We show here only reactions which occur in both models, and which showed convergence across 4 repeated BayFlux runs (*r*^ < 1.10, Gelman-Rubin statistic [38], see main text). Reaction names correspond to Core Metabolic Model 2 (see Materials and Methods).

Interestingly, whereas flux distributions for both genome-scale models and core models are mostly concordant in their means, there are a few instances in which flux measurements present very different averages (Fig. 4). For example, PPCK (phosphoenolpyruvate carboxykinase), PPC (phosphoenolpyruvate carboxylase), and GLUDY (Glutamate dehydrogenase) offer very different flux estimates depending on which model we use (lower values for the genome-scale model). Among those are PPC and PPCK, that catalyze opposite reactions, forming a cycle/cyclic flux, so it is only the net flux that is biologically meaningful, and the net flux is approximately the same (Fig. 4). GLUDY, however, catalyzes the conversion of 2-oxoglutarate into glutamate and is negative for the genome-scale model, but hovers around zero for the core model. This example advises caution in assuming strong inferences from ^13^C MFA since the results may depend significantly on the model used. We suggest that the most consistent and repeatable way to proceed is to derive metabolic models systematically from the genome.

In any case, we can see that the idea of narrowly constrainining all fluxes is, often, misleading. For example, for the cases of F6PA, DHAPT and PYK, the genome-scale model shows very flat probability distributions over large ranges of flux values (Fig. 4).

### Monte Carlo sampling enables probabilistic knockout predictions that are more accurate

By leveraging the full probability distribution provided by BayFlux, we developed and evaluated two novel methods to predict fluxes after a knockout (Fig. 5): Probabilistic ^13^C Minimum of Metabolic Adjustment (P-^13^C MOMA), and Probabilistic ^13^C Regulatory On/Off Minimization (P-^13^C ROOM). Unlike classical MOMA [39] and ROOM [40] (or their ^13^C versions [15]), this approach yields a predicted distribution of flux profiles after a knockout that aims to capture the uncertainty inherent in the initial wild type (WT) flux distribution, and represent that in the prediction. P-^13^C MOMA and P-^13^C ROOM work by first computing the WT flux profile distribution using BayFlux (base flux profile distribution), then choosing a representative set of flux profiles through subsampling, and finally computing the MOMA [39] and ROOM [40] knockout prediction for each flux profile in this set (Fig. S4). We performed this analysis with the data previously published in Toya *et al.* [35] for wild type and two gene knockouts (*pyk* and *pgi*) using a genome scale *E. coli* ^13^C model (see *E. coli* genome-scale model below). To compare the P-^13^C MOMA and P-^13^C ROOM knockout predictions with those obtained through traditional MOMA and ROOM, we used Flux Balance Analysis (FBA) on the same genome scale model used with BayFlux to obtain the base flux profile (maximizing growth after removing the growth rate constraint, but keeping extracellular exchange constraints). We then applied MOMA and ROOM to predict the knockout fluxes for *pyk5h* and *pgi16h* knockouts by assuming all probability is concentrated in a single point (a Dirac delta function) to transform this single flux profile into a probability distribution with the same number of samples used for P-^13^C MOMA and P-^13^C ROOM. In this way, we were able to compute a distance to the experimentally measured fluxes for both knockouts (Fig. 6).

**Fig 5.**
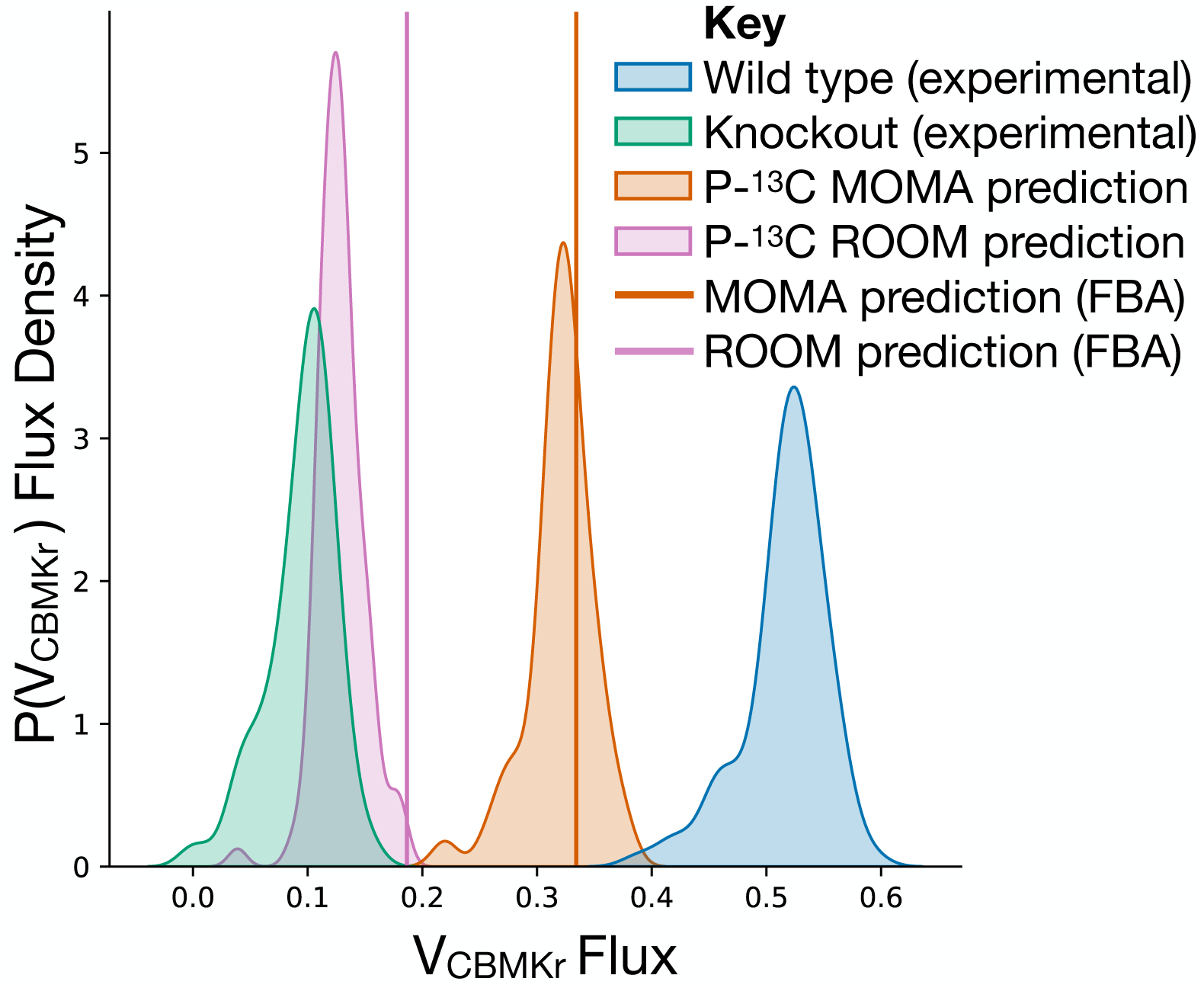
Using BayFlux to predict gene knockout flux distributions improves predictions. We employ P-^13^C MOMA and P-^13^C ROOM (brown and violet respectively) to predict the flux distributions for the *pgi* gene knockout (green, Toya 2010 pgi16h data [35], computed using BayFlux) by leveraging the wild type flux distribution (blue, Toya 2010 wt5h data [35], computed using BayFlux), and compare to FBA based MOMA and ROOM methods (brown and violet vertical lines respectively). Here, we show the reaction CBMKr (Carbamate kinase) as a representative reaction, whereas the distances between full flux profile distributions are provided in Fig. 6. Notice how the probability distributions from P-^13^C MOMA and P-^13^C ROOM are closer to the experimentally obtained probability distribution for the knockout (computed from experimental data using BayFlux), indicating more accurate predictions.

**Fig 6.**
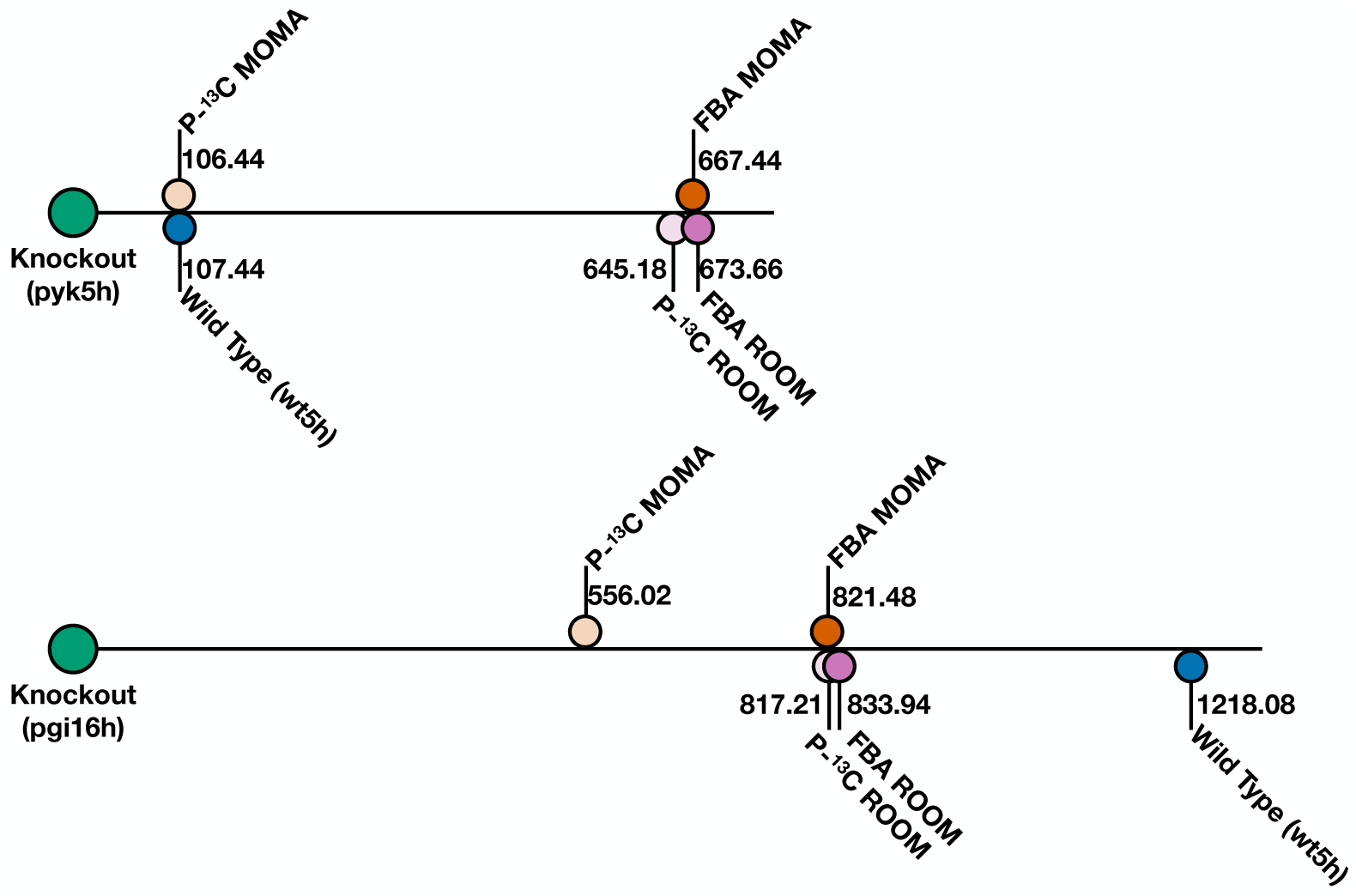
Knockout predictions improve by leveraging BayFlux flux probability distributions. Knockout prediction performance for four methods as judged by the distance of the prediction to the experimentally measured flux profile distribution, as computed with BayFlux from ^13^C experimental data. Distances between flux profile distributions were calculated through a classic measure of how two probability distributions differ from each other: the multivariate Kullback-Leibler divergence [41] (higher value worse prediction, lower value better prediction). Distance between WT base profile distribution and KO experimentally observed flux profile distribution is provided for reference. Notice how P-^13^C MOMA and P-^13^C ROOM produce smaller distances to the experimental results as compared with MOMA and ROOM, indicating improved predictions. All distances are shown on a one dimensional plot as distances from the knockout strains, and do not reflect relative distances between predictions.

The results suggest that knockout predictions improve by leveraging BayFlux flux probability distributions as the base flux distribution for MOMA and ROOM (Fig. 5, Fig. 6). For both knockouts, P-^13^C MOMA outperforms MOMA and P-^13^C ROOM outperforms ROOM. The best predictions for all methods are provided by P-^13^C MOMA, in terms of minimal distance from experimentally measured fluxes (Fig. 6). The code for this analysis is distributed in a Jupyter Notebook along with BayFlux, so it can easily be applied to new knockout prediction problems.

### Evaluation of convergence and scaling performance shows that faster or more efficient sampling will be required for very large systems

The number of required samples to reach convergence using BayFlux seems to scale linearly with the number of reactions in the model (Fig. 7). Convergence, *i.e.* a stable distribution for the flux probabilities, is here defined as at least 80% of achieving a net flux Gelman-Rubin statistic *r*^ < 1.10 [38] (the difference between estimated variance within the same chain and between different chains), across 4 independently run sampler chains. We tested three models: the toy model from Antoniewicz *et al.* [42] (5 reactions), the example *E. coli* core metabolism model from 13CFLUX2 [19] (66 reactions), and the imEco726 genome scale model used throughout this paper [14] (487 reactions with statistical variability, *e.g.* not fixed by stoichiometry). This last, genome-scale, model took 5.86 days to reach convergence after ≈ 33*M* samples using two Intel Xeon Gold 6154 (3 to 3.7 Ghz) cpu cores per sampler, for a sampler performance of approximately 65.44945 samples per second.

**Fig 7.**
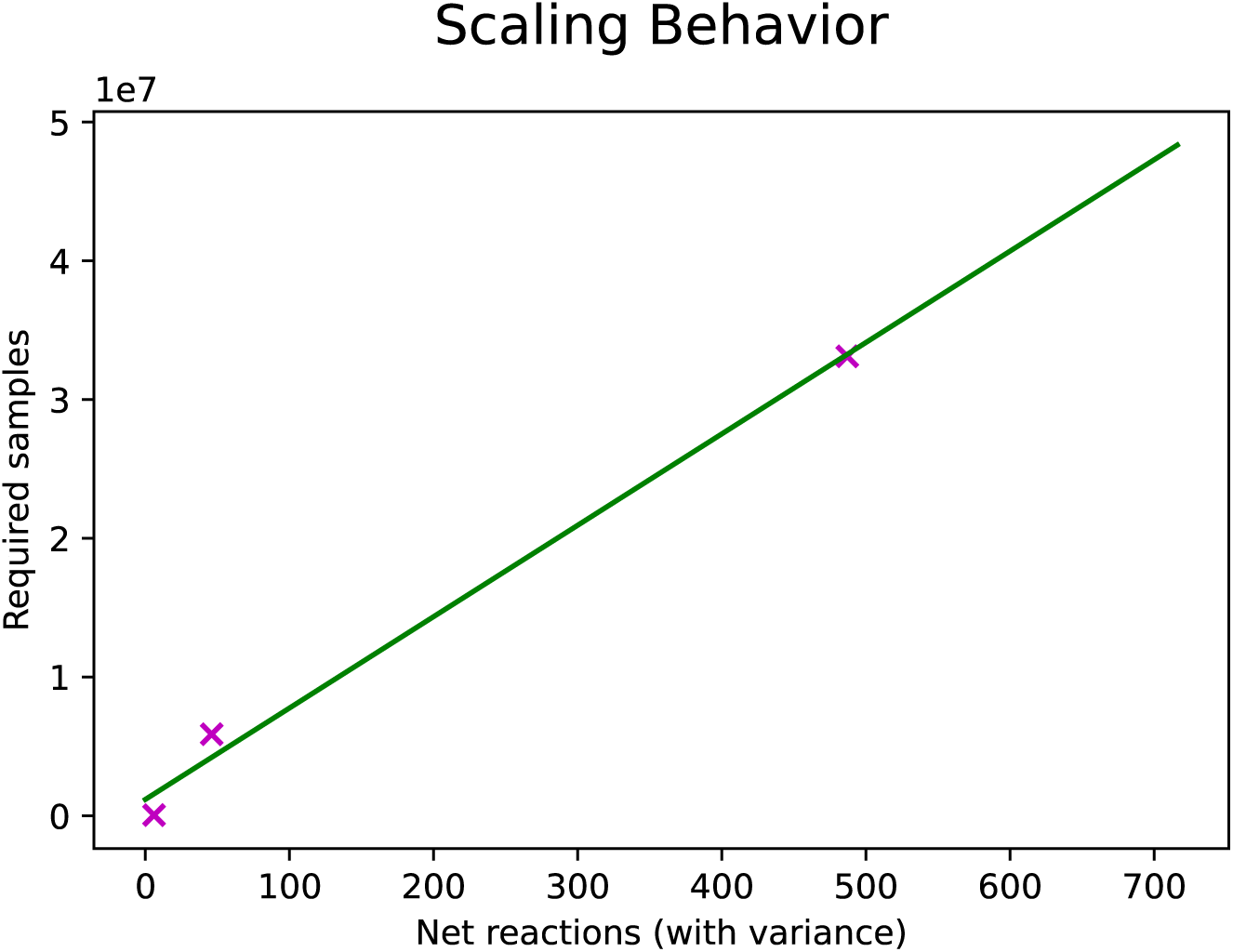
The number of samples needed for BayFlux convergence scales approximately linearly with number of reactions in the model. Shown are the number of samples required to reach convergence across four parallel chains, for three different size models and the fit to a linear model. We define convergence as having at least 80% of reactions with a net flux Gelman-Rubin statistic *r*^ < 1.10 across 4 parallel chains, and exclude reactions with no sampling variance, *e.g.* reactions that are fully constrained and have only a single possible flux value (allowing for small amounts of numerical error) [38]. The data used here for the two largest models are the wild type 5 hour data from Toya *et. al.* [35].

Novel, faster, sampling algorithms will be required to apply BayFlux to the large metabolic models involved in microbial communities and human metabolism. A quick back-of-the-envelope calculation shows that a community of ≈ 200 species (3000 200 = 600, 000 reactions) would require over 19 years to achieve convergence based on the linear slope (65, 881) and intercept (1, 174, 530) shown in Fig. 7, assuming the time per sample were roughly the same 65.44945 samples per second as in our genome-scale *E. coli* model (a reasonable approximation because larger sparser matrices present in larger models are fundamentally better suited to parallelization, making the per-core runtime similar despite increased computing demands):

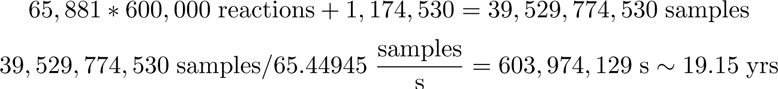

Similarly the recent human metabolic model by Tiele *et al* [43] (80, 000 reactions) would require over 2.5 years to converge:

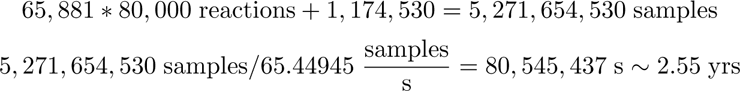

Hence, if we are to tackle these problems in a reasonable amount of time, parallelization or more efficient sampling algorithms are required. BayFlux may be particularly informative for these applications since the best fits to the data produced with optimization are likely to be misleading.

Preliminary tests have shown that we can get approximately an order of magnitude speedup just by using sparse matrix solvers. We have found that we can get a ≈ 8.10*x* speedup by using the popular sparse matrix solver algorithm SuperLU in place of the Numpy dense matrix solver from the Intel Distribution for Python that we are currently using, based on the mean of seven repeated matrix solving steps in our *E. coli* genome scale model [44]. We intend to support this alternative solver in future versions of BayFlux, after incorporating sparse matrix representations into the software.

## Conclusion

We have presented here a method (BayFlux) that provides all fluxes compatible with ^13^C experimental data for a genome-scale metabolic model (Fig. 1). BayFlux works by combining Bayesian inference and Markov Chain Monte Carlo (MCMC) sampling (Figs. 1,8), and produces results compatible with the traditional optimization approach to estimating fluxes through ^13^C MFA (Fig. 2). However, BayFlux results provide extra information in the form of the full flux probability distribution, which allows for a more nuanced understanding of which flux values are possible given the current experimental data set (Fig. 3). Moreover, BayFlux’s rigorous quantification of uncertainty shows that optimization models can overestimate flux uncertainty by representing it through only two numbers: the upper and lower confidence intervals (Fig. 3).

**Fig 8.**
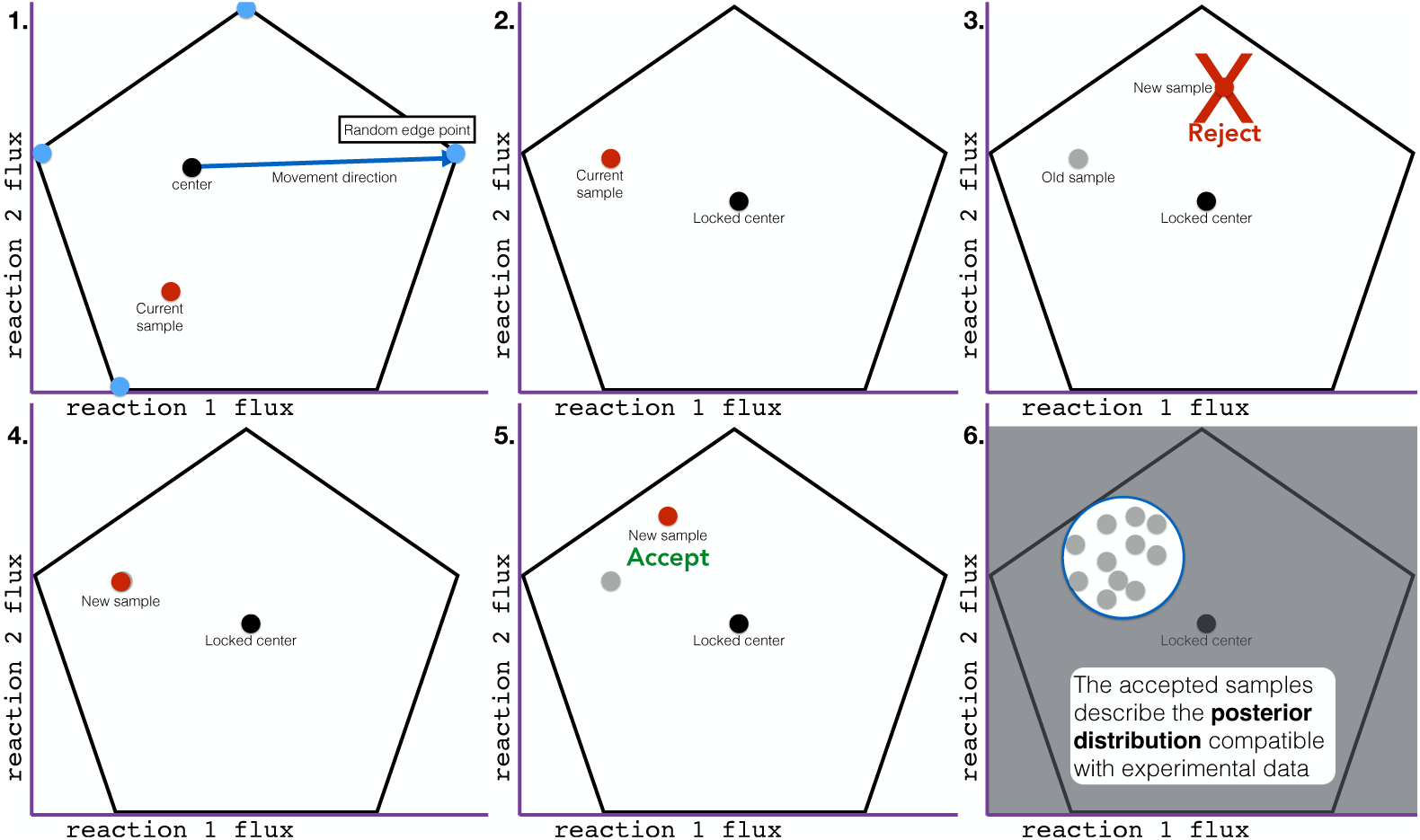
Graphical illustration of Artificial centering Metropolis sampling (AcMet) behind the BayFlux software package. The AcMet algorithm is used to sample the phase space and find the probability for each flux profile (see Eq. (4), and Algorithm 1). Each frame illustrates a step in the AcMet algorithm, shown in only two dimensions for simplicity. The black outline represents the feasible flux polytope, as determined by the genome scale model stoichiometric matrix. **1. Center identification.** Initial ‘edge points’ are identified on the edges of the flux space by minimizing and maximizing each reaction. A running average of all samples is maintained as the ‘center’ and a series of samples are taken, always moving the current point in a direction determined by the current center and one of the edge points. Once a direction is determined, a sample is chosen from the uniform distribution within the allowable bounds, and all samples are accepted. Sufficient samples are collected to obtain a stable center. **2. Metropolis sampling.** Once a stable center is identified, the center is locked, and all previous samples are discarded. New proposed samples are collected in the same manner as step 1., but without updating the center. **3. Reject low probability samples.** Samples are accepted or rejected probabilistically based on the ratio of the likelihood of the data given the new sample, divided by the likelihood of the data given the current sample, *L*(*data new sample*)*/L*(*data current sample*). All higher likelihood samples are accepted. **4. If a sample is rejected, back up a step.** If a sample is rejected, it is discarded, and the sampler is moved back to the previous sample location, and records an additional sample at the previous location. **5. If a sample is accepted, continue.** If a sample is accepted, it is recorded and more samples are collected, just as in step 1, but without updating the center. **6. Halt and report posterior probability.** After a sufficient number of samples are collected, they are used to describe the posterior probability distribution.

Surprisingly, the genome-scale model of metabolism produces much narrower flux distributions than the small core metabolic models that are traditionally used in ^13^C MFA (Fig. 4). We believe that the cause of this increased precision in flux determination for more complex models despite the apparent increase in degrees of freedom is due to the bow tie structure of metabolism [37]. Due to this structure, central metabolic fluxes are well constrained by metabolite labeling, and the inclusion of several hundreds of extra reactions only adds draws of cofactors that further constrain core fluxes. An additional surprising finding is that whereas flux distributions for both genome-scale models and core models are mostly concordant in their means, there are a few instances in which flux measurements present very different averages (Fig. 4). This finding advises caution in assuming strong inferences from ^13^C MFA since the results may depend significantly on the model used. We believe that the most systematic way to obtain a model for ^13^C MFA is to derive it methodically from the genome annotation, as we have done here, by creating a genome-scale model from the genome annotation and augmenting it with the corresponding atom transitions for each reaction.

Based on BayFlux, we developed and evaluated novel methods (P-^13^C MOMA and ROOM) to predict the biological results of a gene knockout, that improve on traditional FBA-based MOMA and ROOM methods (Fig. 5). P-^13^C MOMA and P-^13^C ROOM leverage the full flux probability distributions measured through BayFlux to provide probability distributions of fluxes after a gene knockout in a way that captures the uncertainty inherent in the initial flux. These probability distributions show a lower average distance to the experimentally determined results for the knockout than those provided by the FBA-based methods (Fig. 6).

The scaling properties for BayFlux with respect to model size (Fig. 7) indicate that significant efforts in improving sampling and parallelization will be required to apply this method to large models such as microbiome or human metabolism models (displaying 80,000-200,000 reactions). However, preliminary results suggest that upgrades, such as a sparse solver, can increase speed by orders of magnitude.

In summary, BayFlux provides a rigorous way to find all flux profiles compatible with a given set of ^13^C experimental data, opening the door to an improved understanding of metabolism and more effective predictions for strain metabolic engineering.

## Materials and methods

### Overview of physical problem

The overarching goal of this work is to leverage diverse and noisy experimental data to infer the metabolic fluxes in a cell: *e.g.* the number of chemical species per unit time through every chemical reaction in a living organism. A single cell can encompass thousands to tens of thousands of chemical reactions. We divide these reactions into internal, and external (*i.e.* exchange) reactions, where at least one product or reactant is extracellular. Exchange reactions can be observed directly, by measuring the rate of change in extracellular concentration for a given chemical species: *e.g.* the exchange flux for glucose will match the rate at which the glucose concentration falls in the culture media. Internal or intracellular fluxes can only be inferred indirectly from other types of experimental data, as the chemical species they consume and produce are immediately also consumed and produced by other reactions. For example, a reaction ‘A’ may have a very high flux, but its chemical product can still be almost absent from a cell if reaction ‘B’ consumes its product at the same or a higher rate.

Metabolic flux is a coarse-grained concept, comprising a large number of heterogeneous chemical reactions in a living cell. For very simple systems, it is possible to treat individual models, atoms, and enzymes as distinct events, *e.g.* using stochastic chemical kinetics [45]. However, these types of kinetic simulations are infeasible for a full cell. When a large number of cells are growing exponentially under stable environmental conditions (*e.g.* constant cellular doubling time) the random effects of individual chemical reaction events are cancelled out, allowing us to apply a steady state approximation. This assumes that the quantity of each chemical species has reached a quasi-stable equilibrium, and therefore one can regard each reaction as having a specific flux, where the total flux of reactions generating a chemical species always equals the sum of fluxes consuming it. In addition, applying the simplifying approximation that each cellular compartment or organelle is ‘well mixed,’ allows a single flux value to represent each reaction in each compartment.

### Key challenges

An important challenge involves integrating together diverse data sources with fundamentally different types of error, and underlying relationships to metabolic flux. Most common among these data types are exchange fluxes and mass isotopomer distributions (MIDs). Extracellular exchange fluxes account for mass entering and exiting the cell, and include measurements such as nutrient consumption and metabolic byproduct production rate, as estimated by a time series of extracellular concentration measurements and biomass or growth rate measurements. Exchange fluxes are represented directly as metabolic fluxes for specific reactions, and can be either applied as stoichiometric constraints to specific reactions, or as probablistic constraints on those fluxes. Mass isotopomer distributions (MIDs) are the result of feeding the organism with defined isotopomer nutrient substrates: *e.g.* glucose with extra neutrons on specific atoms. Since different metabolic flux vectors result in shuffling of atoms in different manners, they produce distinct MIDs. Using the Elementary Metabolic Unit (EMU) method [42], we can calculate the MID for any given flux vector, and compare this to the experimental data.

Perhaps the most important challenge is that, given the incredible complexity of metabolism and the paucity of (noisy) experimental data, the metabolic flux system is severely underdetermined, with substantial uncertainty about the true metabolic flux profile. As shown in the examples below, the number of fluxes for a genome-scale model is typically on the order of *∼* 3000, whereas the available data to constrain is usually on the order of 1-5 exchange fluxes and *∼* 60 metabolite measurements.

### Mathematical problem formulation

Mathematically, we represent metabolism as a directed bipartite graph, with *m* metabolites and *n* reaction vertices. Metabolites represent unique chemical species in distinct compartments or areas of the organism. They participate as reactants in chemical reactions, and produce new chemical species, *e.g.* products, as a result. Reactants are represented as directed edges from the metabolite vertices to a reaction vertex, and products are represented as directed edges from the reaction vertices, to new metabolite vertices. This is the standard method used for genome scale models, as it encodes the chemical stoichiometry information for each reaction.

We represent the metabolic flux through this system with the unknown vector of fluxes **v** *∈* R*^n^* associated to each reaction, with units of molecules per cell biomass unit per time. For example, the flux *v_P DH_* = 0.16 mM/gDW/h represents that the pyruvate dehydrogenase reaction is converting 0.16 mM of pyruvate per gram of cell dry weight per hour into Acetyl-CoA. Our system is constrained by conservation of mass, as described by the stoichiometric matrix *S* where:

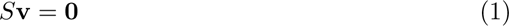

at steady state [46], enforcing that the mass consumed by each reaction matches the mass produced.

Our goal is to characterize fluxes probabilistically, *i.e.* find the joint distribution of fluxes given data:

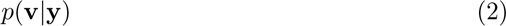

where **y** represents the experimental data:

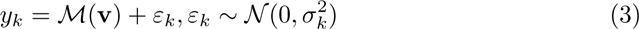

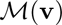 is the function of the simulated MIDs (details in the “Software Implementation” section below), and (optionally) any other relevant experimental data. For the MID data, we assume a Gaussian error model with a zero mean and standard deviation *σ_k_* for each measurement *k* in the likelihood function.

For the purpose of finding the joint distribution, we use a Bayesian inference approach [47]):

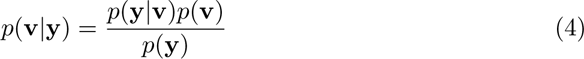

The likelihood function follows from Eq. (3), assuming that measurements are i.i.d, and normally distributed:

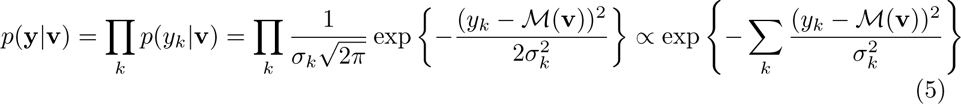

For simplicity, we use a uniform prior *p*(**v**) (*i.e.* a priori probability that the flux vector is **v**) on the polytope *S* defined by *S***v** = 0: *i.e.* 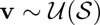. However, it is straightforward to modify this to incorporate any additional prior knowledge. When extracellular exchange fluxes are measured with high accuracy they can be added directly as stoichiometric constraints such that the prior is a uniform probability distribution on the set 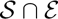, where *ε* is defined by experimentally measured exchange fluxes, and zero probability outside. Alternatively, if there is substantial experimental uncertainty, the extracellular exchange fluxes could be added to the likelihood function.

The marginal likelihood (normalizing constant in Eq.(4)) 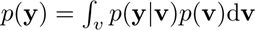 is difficult to compute, is only known for a small class of distributions, and becomes intractable with a high number of dimensions [48]. Therefore, we can only compute *p*(**v|y**) up to a normalizing constant, *i.e.* 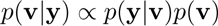, for which we use Markov Chain Monte Carlo (MCMC). MCMC allows us to draw samples from *p*(**v|y**) by creating a Markov chain that converges to the target distribution *p*(**v|y**) [48]. Our proposals are generated by choosing a random direction, and jumping along that direction to a uniformly distributed sample within the stoichiometric bounds on *S* (Fig. 8). A generated proposal **v**^′^ is accepted according to the Metropolis probability [48, 49]:

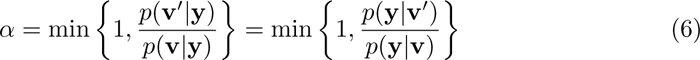

The normalizing constant cancels out, and so does the prior, since it is uniform. In the current implementation we use the AcMet algorithm, described below to compute the proposal.

### Sampling

We created a new algorithm (Artificial centering Metropolis sampling, or AcMet, algorithm 1, Fig. 8) for sampling from the posterior probability distribution, based on the commonly used Artificial Centering Hit-and-Run (ACHR) algorithm [33]. The ACHR algorithm is widely used to collect pseudo-uniform samples from the stoichiometrically feasible flux polytope of genome scale metabolic models (GSMMs). We modified the ACHR algorithm and combined it with the Metropolis algorithm, to produce a Markov Chain Monte Carlo (MCMC) sampler for Bayesian inference of metabolic fluxes (AcMet). AcMet can sample from the posterior probability distribution obtained by updating priors with an array of experimental evidence that includes ^13^C isotopomer mass distribution vectors for metabolites, as well as extracellular exchange fluxes computed from time series measurements of extracellularmetabolite concentrations. Other experimental data can be added in a similar fashion, facilitating integration of diverse omics data into a single coherent model.

The ACHR algorithm presented two characteristics that made it unsuitable for Bayesian inference with ^13^C experimental data: center updates and the need to sample from net fluxes rather than both directional components of reversible reactions. Center updates involve finding the center of the accessible flux volume, which is used to direct the next sampling point [33]. These center updates happen in every step of the ACHR algorithm, and are not a problem in practice when doing uniform sampling of genome scale models. However, this ‘moving center’ makes the sampler non-reversible, eliminating the ability of the MCMC process to sample from the posterior. Moreover, it is critical for efficient sampling that the ‘center’ remains in the true center of the flux polytope. However, when sampling a non-uniform distribution, *e.g.* for Bayesian inference, samples no longer average around the center of the flux polytope, which would make using these samples to compute the center impossible. Therefore, maintaining a fixed center is critical to enable Markov-Chain Monte Carlo sampling. Sampling from net fluxes only is a problem because of cyclic reaction fluxes, which involve simultaneous forward and backwards fluxes for the same reaction [50]. While cyclic reaction fluxes play no role in the stoichiometric FBA simulations that originated the ACHR algorithm, they are critical for determining metabolite labeling in ^13^C MFA [50].

AcMet overcomes the center updates problem by sampling in two phases. First we collect a large number of uniform samples from the genome scale model using the ACHR algorithm, until the center converges into a stable position. Next, we lock the center while collecting samples with the Metropolis algorithm, which accepts or rejects samples taking into account the likelihood function derived from experimental data. By keeping the center locked, the chain becomes reversible, permitting Metropolis sampling. The ACHR algorithm begins with finding ‘edge points’ (see Glossary of terms) that are coordinates on the most extreme bounds of the feasible flux space, which are points the sampler can move towards, and are used to navigate around the unusual shape of the flux polytope. Typically these represent the highest and lowest points in each dimension (*e.g.* reaction flux), as identified by linear optimization. Here, we use the term ‘edge points’ instead of the more commonly used term ‘warm-up points’ to avoid confusion with the Markov Chain Monte Carlo (MCMC) concept of warm-up samples. These ‘edge points’ are coordinates in the feasible flux space, and should not be confused with the concept of edges in a mathematical graph.

In order to effectively sample within-reaction cyclic fluxes, we added extra edge points to the sampler, allowing the sampler to explore along these dimensions, in addition to exploring net fluxes. For each reversible reaction, two additional edge points were added: one which maximizes forward flux and minimizes reverse flux, and another which minimizes forward flux and maximizes reverse flux, as computed from the allowable bounds of each reaction.

#### Algorithm 1 Artificial centering Metropolis sampling (AcMet, see Fig.8 and Table 1).

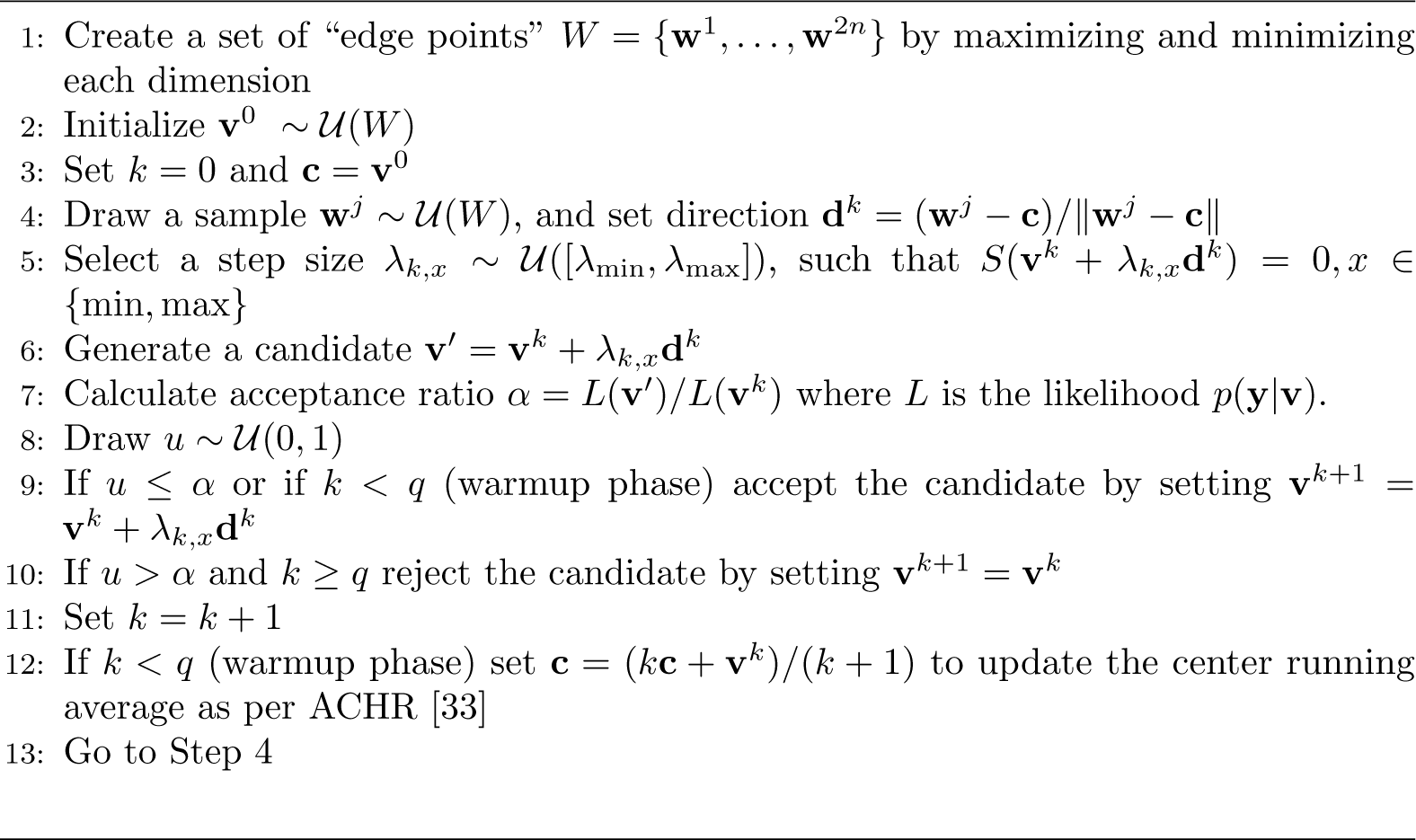

**Table 1.** Notation.

| Variable | Description |
| --- | --- |
| $n$ | Number of reactions |
| $m$ | Number of metabolites |
| $\mathbf{v}$ | $n$-dimensional vector of fluxes |
| $\mathbf{c}$ | The center of the flux polytope (*e.g.* a running average of uniform samples) |
| $q$ | The number of quasi-uniform samples collected before locking the center |
| $\mathcal{U}(\cdot)$ | Uniform probability distribution |
| $W$ | The set of “edge points” |
| $k$ | Flux sample counter |
| $x$ | The set of all valid proposals within the flux polytope from the current sample in a given direction |
| $u$ | A random sample collected from the uniform distribution |
| $\mathbf{d}^k$ | The proposal direction for a new sample |
| $S$ | Stoichiometric matrix representing the products and reactants for each chemical reaction |
| $\mathbf{y}$ | A vector representing all experimental data used to infer metabolic flux |
| $y_k$ | Experimental data for observation $k$ |
| $\sigma_k$ | Standard deviation of error for observation $k$ |
| $\varepsilon_k$ | Experimental error for observation $k$ |
| $\mathcal{E}$ | The convex polytope representing valid fluxes, based on measurement of extra-cellular exchange fluxes |
| $\mathcal{S}$ | The convex polytope representing valid fluxes |

The likelihood *L* is computed for each step according to the normally distributed likelihood function (Eq. (5)). A uniform prior is implicitly included when calculating the acceptance ratio (eq. (6)).

### Models for generating metabolite labeling

#### Core metabolic model 1

To compare our method with existing ^13^C MFA software, we used the *E. coli* data and core metabolic model employed in the demonstration of 13CFLUX2 (the slightly adapted version of [51] used in [34]). The experimental data involve measurements of glucose uptake, growth rate, and the labeling of eleven central carbon intracellular metabolites for an *E. coli* MG1655 strain grown in glucose-limited continuous culture. The model comprised 66 reactions and 37 metabolites describing central carbon metabolism (selected reactions are shown in Fig. 2). In order to provide enough instances for a comparison, we randomized the exchange fluxes fifteen times and determined fluxes through both 13CFLUX2 and BayFlux (Fig. S2).

#### Core metabolic model 2

To compare genome scale and core metabolic models, we used the previously published Toya 2010 wild type 5 hour (wt5h) data, which includes measurements of glucose uptake, acetate excretion, growth rate, and the labeling of nine central carbon intracellular metabolites for an *E. coli* BW25113 strain [35]. As the core metabolic model, we used a simplified *E. coli* core model with 63 reactions taken from previous literature [15], and as the genome scale model the *E. coli* genome scale model described in this paper which combines iAF1260 and imEco726 models. Note that both models have a built in biomass composition, which we left intact as originally described in each model, and are not directly comparable because they merge and separate different metabolites.

#### Genome-scale *E. coli* atom mapping model

To evaluate the method described by this paper using real world data, with a genome scale model, we merged a genome scale ^13^C MFA model with a genome scale stoichiometric model, and used this to evaluate previously published *in vivo* ^13^C MFA data.

We utilized atom transitions from the imEco726 genome scale isotope mappings, together with the latest version of the iAF1260 metabolic reconstruction of *Escherichia coli* [14, 36]. We updated both the iAF1260 and imEco726 models to account for the experimental conditions used by Toya *et al.*, and accommodate recent updates to the latest version of iAF1260, including new reactions and changes to some reaction and metabolite identifiers [35].

This process was facilitated by the fact that imEco726 was initially based on iAF1260. Merging these models and adjusting them to be suitable for a specific experimental context required numerous refining steps, as described below. The full source code for this model creation and curation process is included with the BayFlux software package as a Python Jupyter notebook, such that it can serve as a template for users to repeat the general process for other organisms.

##### Mapping metabolite and reaction identifiers

The iAF1260 genome scale model contains 2382 reactions, and the imEco726 isotope mapping model contains carbon transitions for only 686 reactions. We found that 70 of the transitions in imEco726, and all of the metabolites participating in transitions (595 total) could not be mapped to any identical identifiers in the latest version of iAF1260. To address this issue, we performed a text based alignment using the Levenshtein distance between the unmapped identifiers in both models, which we then manually reviewed for correctness [52]. By inspecting the alignment output, we created a set of regular expressions which correctly map all reaction and metabolite identifiers in imEco726 to the corresponding identifiers in iAF1260.

For some reactions with the same or similar names we found that the products and reactants were swapped between the two models. For these reactions, we reoriented the transitions from imEco726 to match those of iAF1260. Also, for this analysis, we are assuming steady state labeling, so we have omitted the “dil” transitions from imEco726, which account for metabolites which have not yet reached steady state ^13^C labeling.

##### Metabolite symmetry

Some metabolites in the genome scale model (*e.g.* succinate) exhibit structural symmetry, such that there is no single unique way of numbering the atoms. For reactions which act on symmetric metabolites, we duplicated the reactions atom transition(s) such that equal flux flows through the model with all possible orientations of the symmetric metabolite. Our software allows for an unlimited number of atom transitions for each reaction, over which the total flux for the reaction is equally divided.

##### Setting flux bounds and extracellular exchange fluxes

After incorporating measured extracellular exchange fluxes for biomass, glucose uptake, and acetate production to iAF1260 we set a maximum absolute value flux for each reaction to 5× the glucose uptake rate, in order to constrain the search space, and accelerate sampler convergence.

##### Applying cell culture media constraints

As an initial step to simplify the model, which contains a large number of infrequently used extracellular exchange reactions, we applied an automated method to pair down these reactions, which we then manually refined for correctness. After incorporating the experimentally measured extracellular exchange fluxes to iAF1260, we computed the smallest set of essential media components using Mixed Integer Programming (MIP) as implemented in the COBRApy function minimal media with the option minimize components set to True [53]. This resulted in 15 essential nutrients that must be included in the media. We applied this minimal media constraint to the model, by setting the lower bound of all non-essential exchange reactions to zero.

Next, to manually refine these for correctness, we obtained the actual experimentally used media composition as published by Toya et al., and identified the set of exchange reactions that correspond to the molecules in this media formulation. From this, we found that two additional nutrients (H_2_O and Na^+^) were provided in the media that were not identified as essential in the analysis described above, and we enabled the uptake of these.

### Model reduction

#### Pruning non-essential reactions with unknown atom transitions

Next, we utilized Parsimonious Flux Balance Analysis (pFBA) to identify non-essential reactions in the iAF1260 model which had also been omitted from imEco726. We found 1655 non-essential intracellular reactions, of which 1310 can both carry carbon and did not have provided transitions in imEco726, so we removed them from the model, along with 499 (now unused) metabolites.

#### Pruning unused reactions

Next, we removed all reactions in iAF1260 which cannot carry flux under the experimentally derived extracellular exchange flux conditions, even if they can carry carbon, or have transitions from imEco726. This removed an additional 380 reactions and 371 metabolites from the model.

#### Inferring unknown atom transitions

After removing unused reactions from the iAF1260 model as described above, we identified 35 reactions which are both essential under our extracellular exchange flux conditions, do carry carbon, but do not contain transitions in imEco726. For these reactions, it was necessary to define atom transitions in order to obtain a complete genome scale isotope mapping model.

As a first approximation for computing atom transitions without using chemical structures, we identified the number of carbons in each metabolite based on the chemical formula, and sorted the reactants and products by increasing carbon count. For reactions where no two reactants have the same carbon count, and both the reactants and products have an identical set of carbon counts per molecule, we assumed direct 1:1 transitions between the identically sized reactants and products.

After applying the assumptions described above to estimate transitions for some reactions, we still had 8 reactions flagged as ambiguous, with either equal sized reactants, or a different distribution of sizes between reactants and products suggesting carbon exchange. For these reactions, we manually looked up the corresponding reaction on MetaCyc, and manually wrote transitions that match the atom ordering used in imEco726.

After all of the steps described above including removing reactions, removing metabolites, and adding transitions our final genome scale isotope mapping model contained 692 reactions, and 798 metabolites. Of these 692 reactions, 633 carry carbon and have one or more atom transition mapping.

Please refer to the Jupyter Notebook entitled “imEco726 genome scale” distributed with our BayFlux software, which outlines the full model curation process described above. All steps can be automatically reproduced, and used as a template for recreating this process with other species and/or experimental conditions.

### Software implementation

#### Simulating metabolite labeling

During the process of Bayesian inference from a metabolic model, our software simulates the mass isotopomer distributions (MIDs) for each flux sample and plugs them into the likelihood equation (Eq. (5)). Because this complex simulation must be performed for each sample, developing a high performance Elementary Metabolic Unit (EMU) simulation method was an essential technology to make our method feasible.

We developed a high performance implementation of the Elementary Metabolic Unit (EMU) method capable of simulating genome scale mass isotopomer distributions for millions of flux vectors in just a few hours on standard computer hardware [42]. Three main techniques were utilized to achieve this performance. First, highly repetitive vector and matrix computations are performed directly via low level Fortran code and calls to the Linear Algebra PACKage (LAPACK) and The Netlib Basic Linear Algebra Subprograms (BLAS). Second, we implement ‘EMU pruning’ where long branch free metabolic pathways are automatically identified, and collapsed into a single EMU transition without loss of information. Third, we implement ‘transition merging’ where functionally equivalent atom transitions (*e.g.* from parallel reactions with identical transitions) are automatically identified and merged, also without loss of information.

This Elementary Metabolic Unit (EMU) code also provides the option of defining extra metabolites to simulate beyond those measured experimentally, *e.g.* to cross validate a model with experimental data by leaving it out and re-inferring it. Additionally, it opens the possibility of generating simulated ^13^C experimental data for experimental design purposes. For example, one can simulate different sets of mass isotopomer distribution (MID) data for a wide array of labeled substrates, and measured metabolites. Next, one can perform Bayesian inference on these simulated data with BayFlux to predict the uncertainty surrounding key reactions of interest, and then select an experimental design that provides maximum information on these specific reactions. Additional metabolites to simulate must be defined while processing a model with BayFlux, because, for performance reasons BayFlux automatically avoids computing portions of a model unnecessary to simulate a given set of experimental data.

#### Software package

We provide BayFlux, an open source implementation of the software and algorithms used for this analysis. BayFlux is a Python library for both genome scale and two-scale ^13^C Metabolic Flux Analysis (MFA). It is built to be used in conjunction with the COBRApy library for Flux Balance Analysis (FBA), and provides compatibility with all COBRApy features for analyzing genome scale models [53]. Our AcMet algorithm is implemented as an extension to the existing metabolic sampler infrastructure included in COBRApy. BayFlux can be downloaded from https://github.com/JBEI/bayflux, and includes extensive documentation, as well as automated testing for correctness (unit testing). The git repository also contains Jupyter notebooks with all of the analysis presented in this paper, which can serve as a template for performing a new analysis with BayFlux. BayFlux also works directly with models pre-processed through the Limit Flux To Core (lftc) software library provided at https://github.com/JBEI/limitfluxtocore [15, 37]. This enables BayFlux to perform 2S-^13^C MFA, a simplifying assumption that reduces compute requirements, and makes model curation easier by only requiring atom transitions for central carbon metabolism.

## Glossary of terms

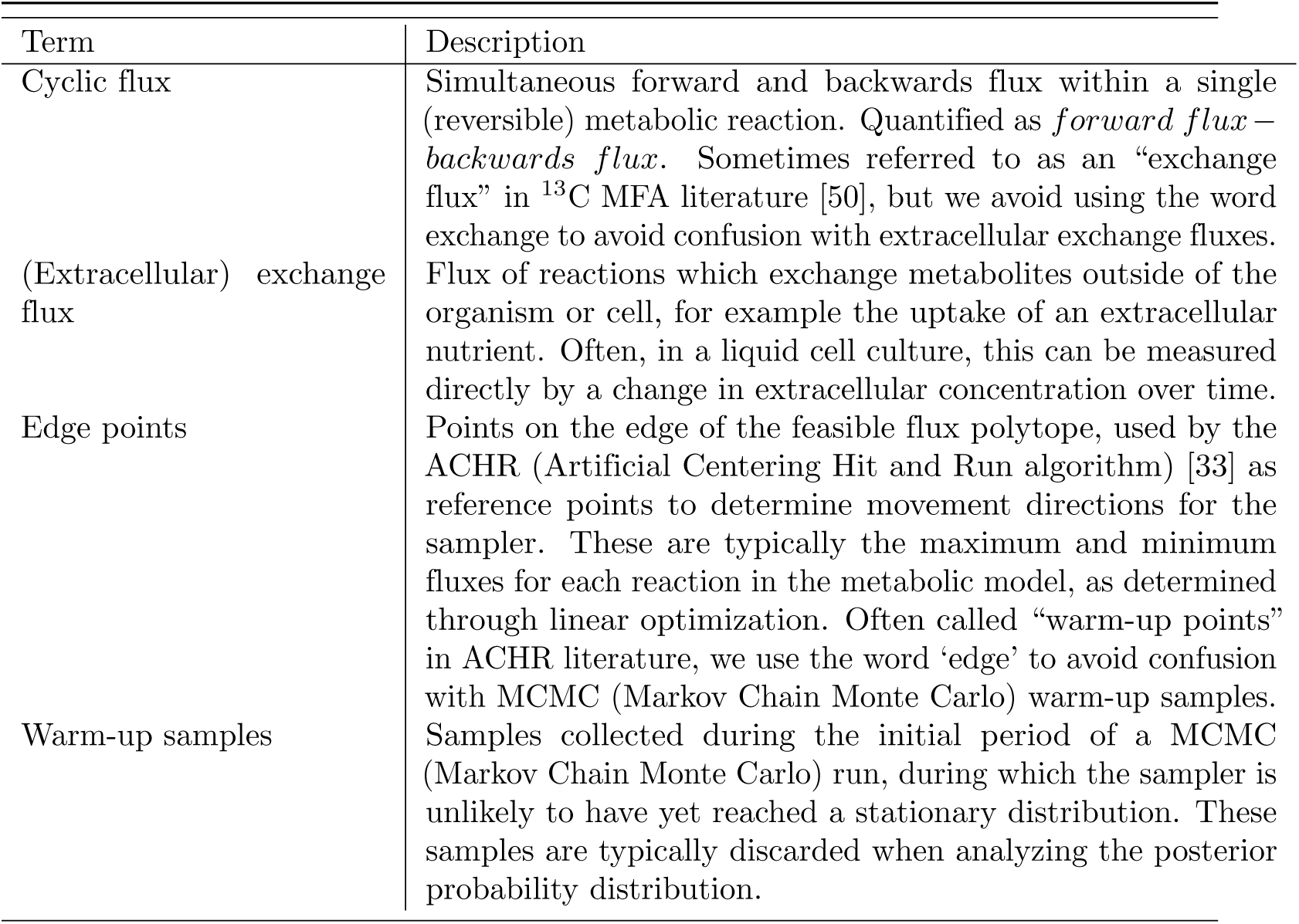

| Term | Description |
| --- | --- |
| Cyclic flux | Simultaneous forward and backwards flux within a single (reversible) metabolic reaction. Quantified as *forward flux* – *backwards flux*. Sometimes referred to as an “exchange flux” in <sup>13</sup>C MFA literature [50], but we avoid using the word exchange to avoid confusion with extracellular exchange fluxes. |
| (Extracellular) exchange flux | Flux of reactions which exchange metabolites outside of the organism or cell, for example the uptake of an extracellular nutrient. Often, in a liquid cell culture, this can be measured directly by a change in extracellular concentration over time. |
| Edge points | Points on the edge of the feasible flux polytope, used by the ACHR (Artificial Centering Hit and Run algorithm) [33] as reference points to determine movement directions for the sampler. These are typically the maximum and minimum fluxes for each reaction in the metabolic model, as determined through linear optimization. Often called “warm-up points” in ACHR literature, we use the word ‘edge’ to avoid confusion with MCMC (Markov Chain Monte Carlo) warm-up samples. |
| Warm-up samples | Samples collected during the initial period of a MCMC (Markov Chain Monte Carlo) run, during which the sampler is unlikely to have yet reached a stationary distribution. These samples are typically discarded when analyzing the posterior probability distribution. |

## Supporting information

S1 File. This file contains supplementary figures.

## Acknowledgements

This work was part of the DOE Agile BioFoundry (http://agilebiofoundry.org), and the DOE Joint BioEnergy Institute (http://www.jbei.org), supported by the U. S. Department of Energy, Bioenergy Technologies Office, and the Office of Science, through contract DE-AC02-05CH11231 between Lawrence Berkeley National Laboratory and the U.S. Department of Energy. This work was also supported by the Basque Government through the BERC 2022–2025 program and by the Spanish Ministry of Science and Innovation MICINN (AEI): BCAM Severo Ochoa excellence accreditation CEX2021-001142-S and PID2019-104927GB-C22 grants.

J.J.C. was supported by the U.S. Department of Energy, Office of Science, Office of Workforce Development for Teachers and Scientists, Office of Science Graduate Student Research (SCGSR) program. The SCGSR program is administered by the Oak Ridge Institute for Science and Education (ORISE) for the DOE. ORISE is managed by Oak Ridge Associated Universities (ORAU) under contract number DE-SC001464.

This research used resources of the National Energy Research Scientific Computing Center, a DOE Office of Science User Facility supported by the Office of Science of the U.S. Department of Energy under Contract No. DE-AC02-05CH11231.

We would like to thank Christian Diener, Niko Sonnenschein, and the rest of the COBRApy team for meaningful discussions, and support in integrating BayFlux with COBRApy. We thank Xiaoye Sherry Li for feedback and meaningful discussions. We would also like to thank Mark Kulawik for providing computer support.

## Author Contributions

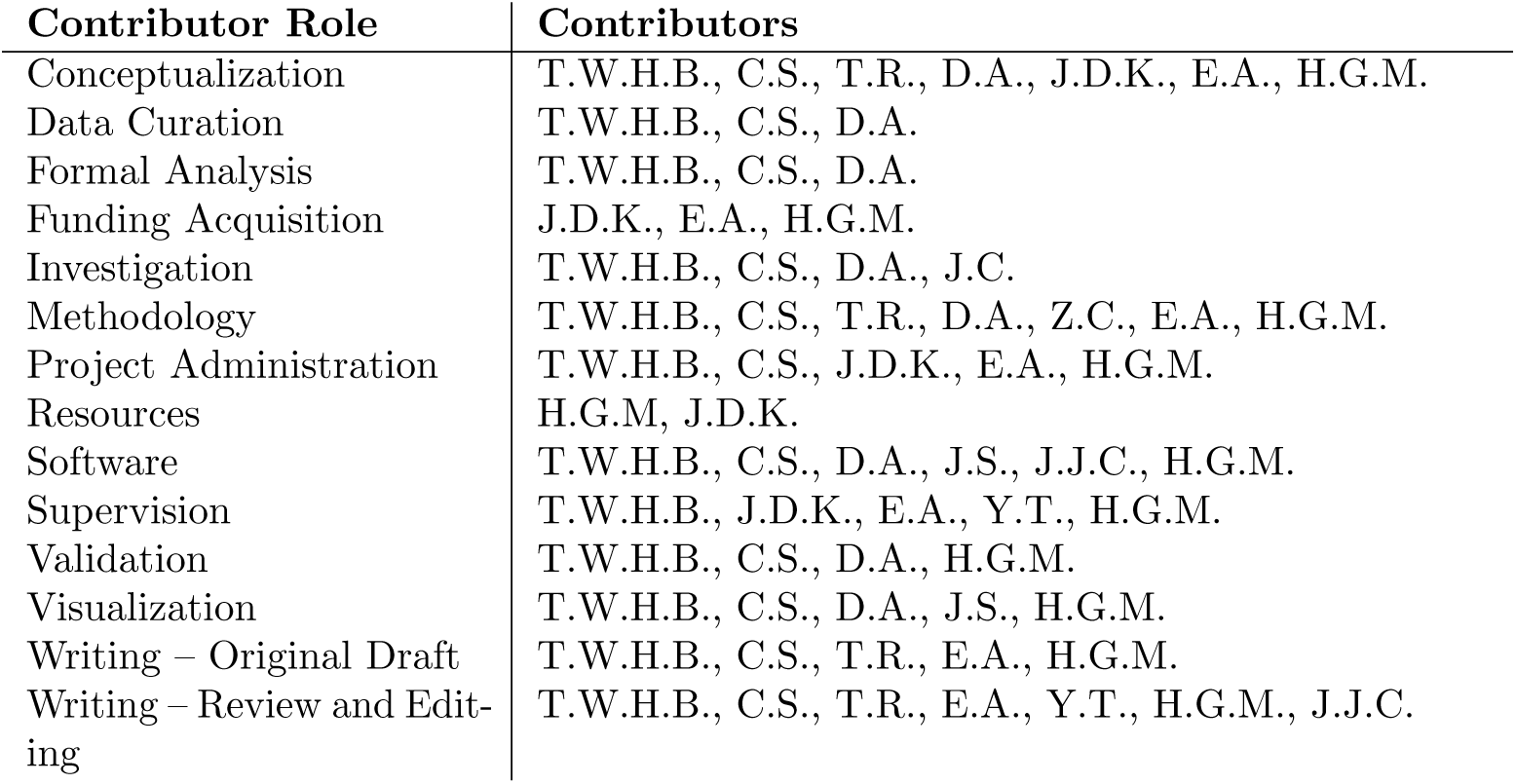

## Supporting Information

**Fig S1.**
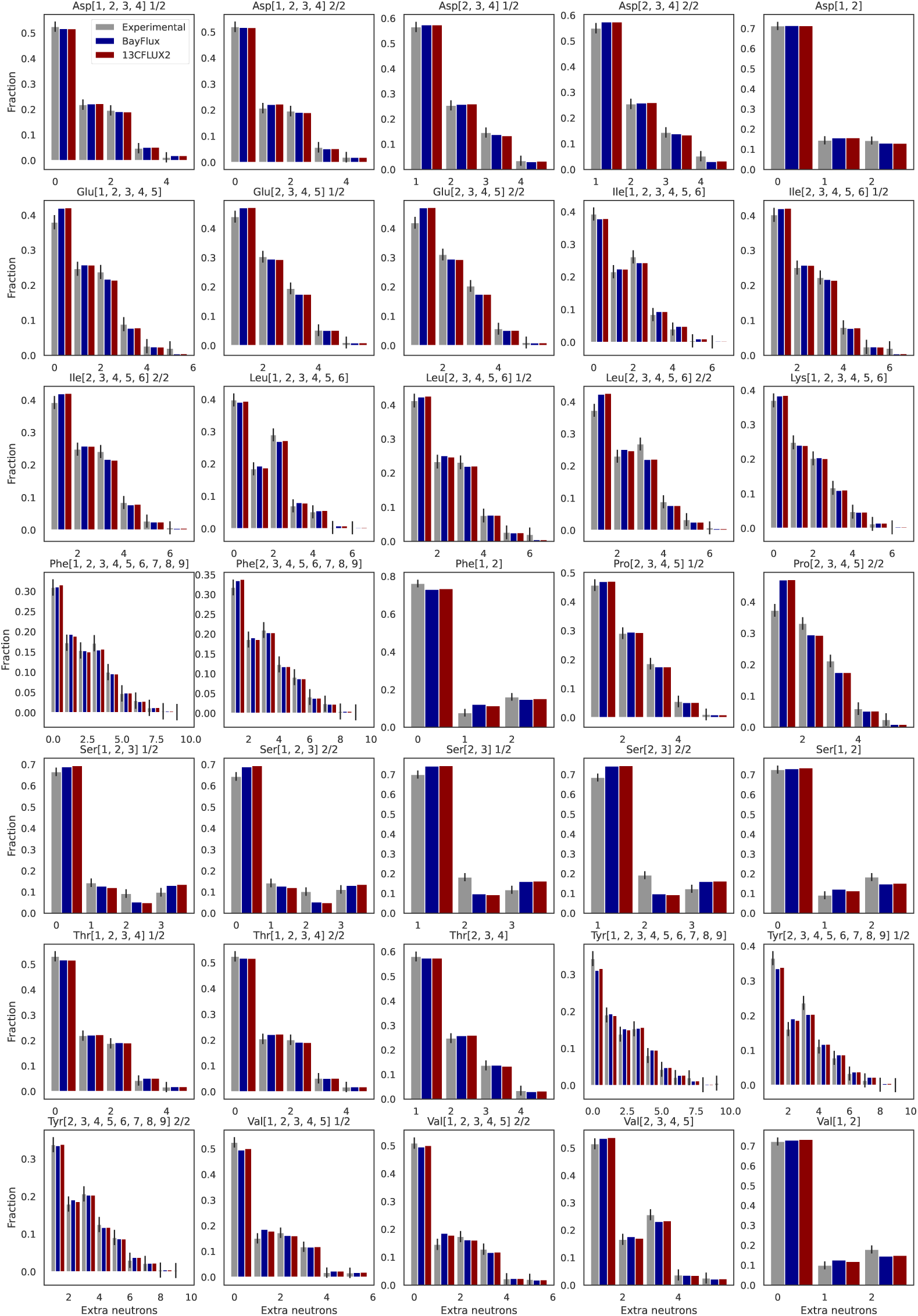
Fit labeling patterns are the same for both approaches, BayFlux and 13CFLUX2. Mass distributions vectors (MDVs) for experimental data (grey), the fits for the best sample for BayFlux (dark blue, ten million samples), and the fits for 13CFLUX2 (red) are closer together than the standard deviation of the experimental error. Units for horizontal axis are the number of extra neutrons in the measured metabolite [54].

**Fig S2.**
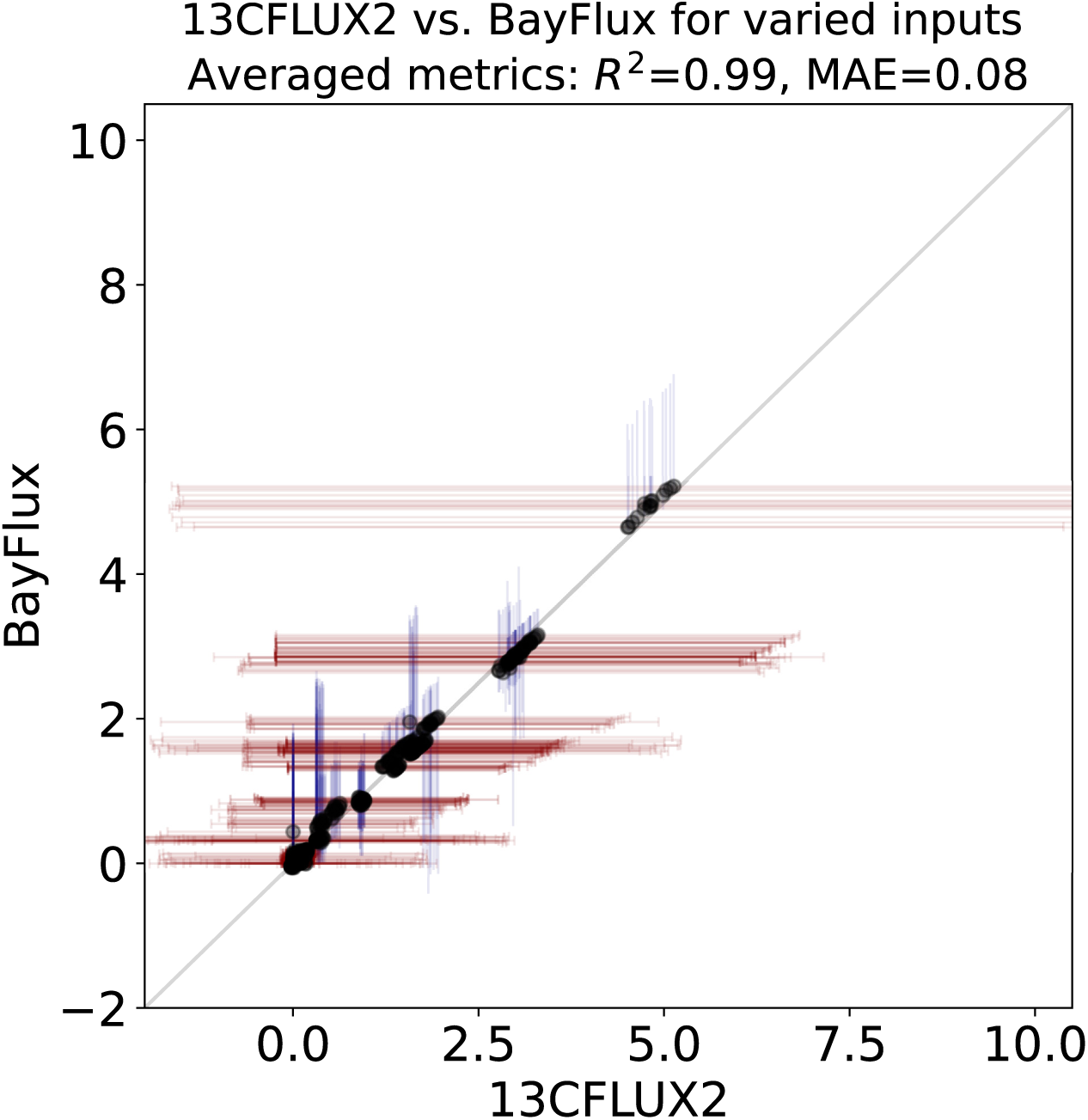
Flux results for BayFlux and 13CFLUX2 are essentially the same for fifteen different inputs. The best BayFlux sample (dark blue, ten million samples, y axis) and its credible interval overlap with the 13CFLUX2 best fit (in red, x axis) and its confidence interval for all fifteen different inputs and fluxes. The fifteen different inputs were obtained by randomly changing the exchange fluxes, but keeping the same labeling profiles for the metabolites. Both axes are in units of mM/gDW/h. We set the 13CFLUX2 confidence bounds that could not be determined to the overall maximum BayFlux bounds for comparison purposes.

**Fig S3.**
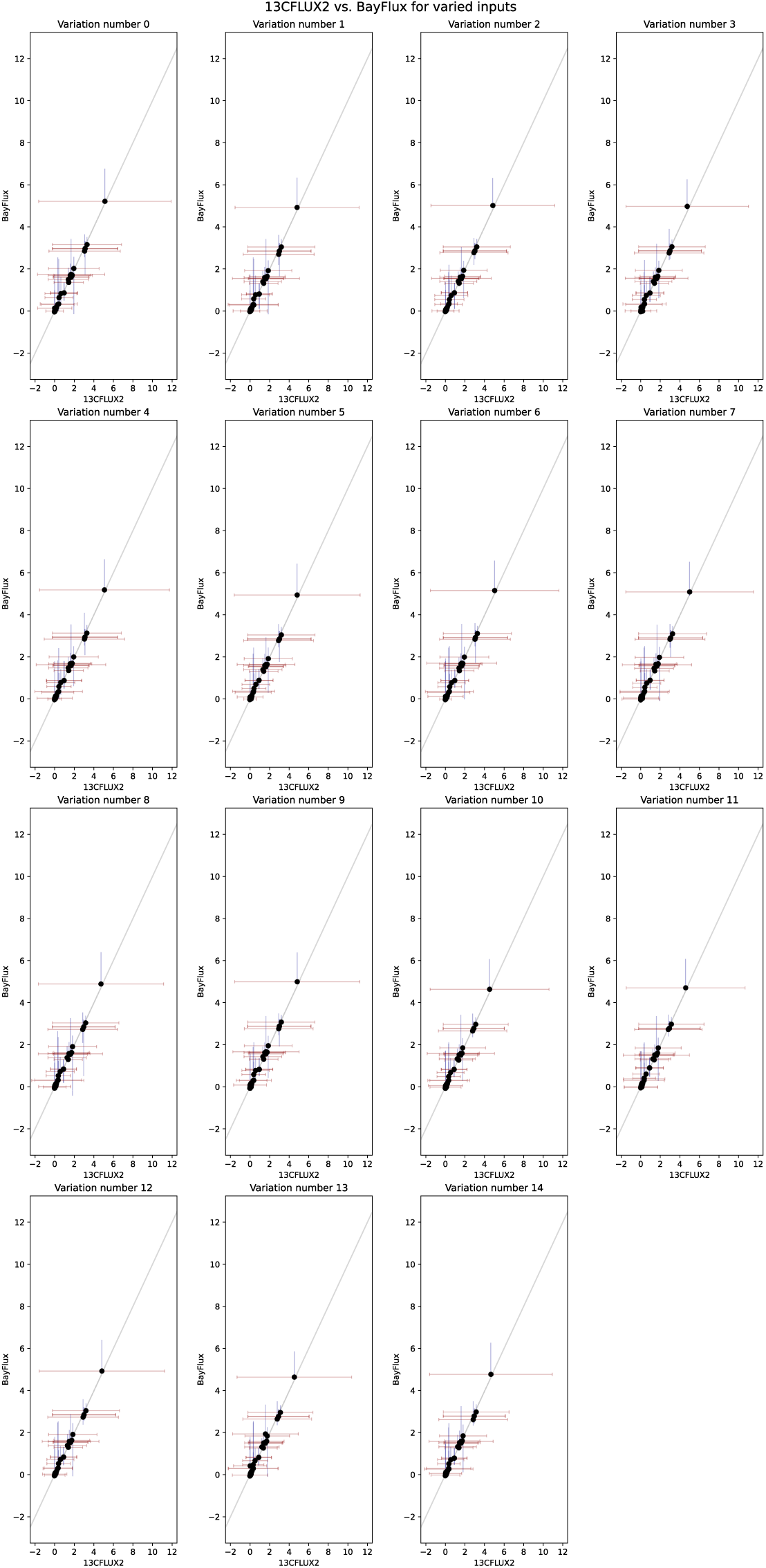
13CFLUX2 vs. BayFlux for 15 different inputs: Best Sample out of 10 million from BayFlux (in dark blue) vs. 13CFLUX2 (in red). We set the 13CFLUX2 confidence bounds that could not be determined to the overall maximum BAYFLUX bounds for comparison purposes.

**Fig S4.**
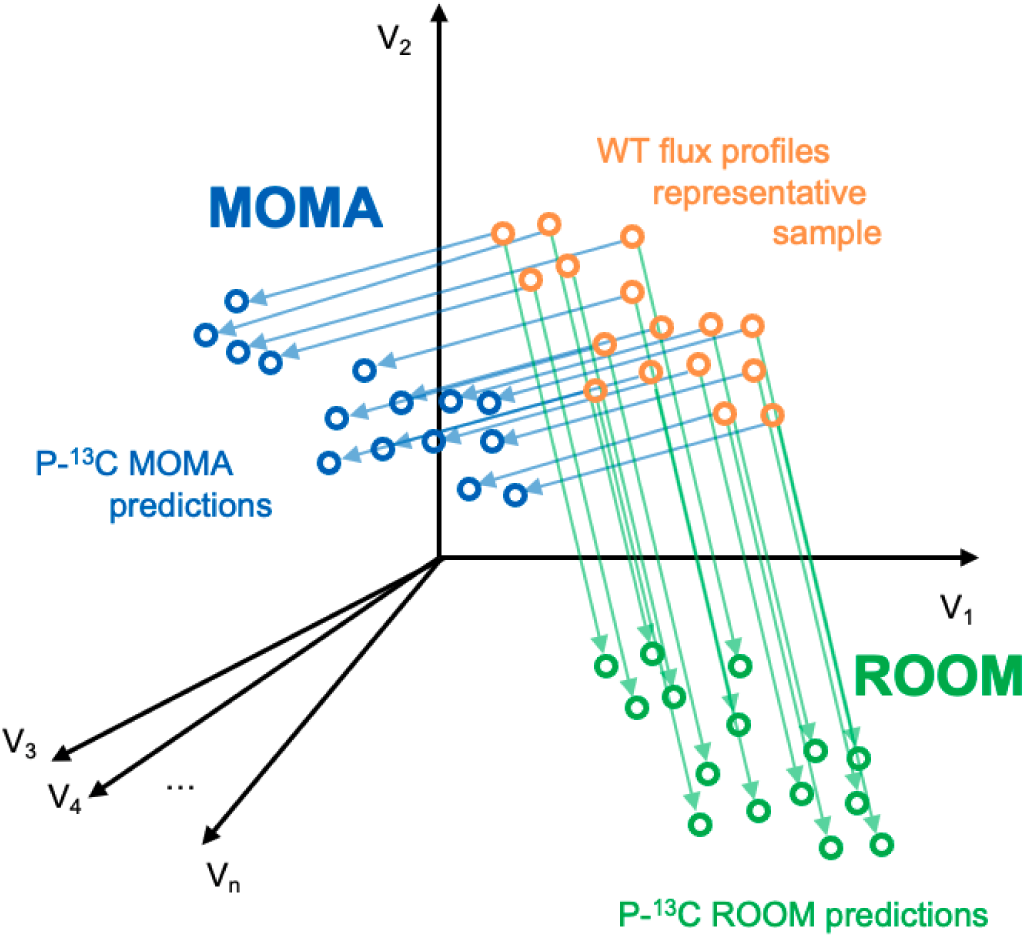
P-^13^C MOMA and P-^13^C ROOM work by first computing the WT flux profile distribution using BayFlux (in orange). Each dot corresponds to a flux profile (*i.e.* reaction fluxes for a full genome-scale model) in the flux phase space. A representative set of flux profiles for this base flux profile distribution is obtained through sampling (orange dots). For each of these flux profiles, a new flux profile is computed by using MOMA (in blue) or ROOM (in green) to predict the resulting flux after a knockout. The fact that BayFlux produces a flux profile distribution results in MOMA and ROOM predicting distributions of flux profiles.

