## Supporting Information for "BayFlux: A *Bay*esian method to quantify metabolic *Flux*es and their uncertainty at the genome scale"

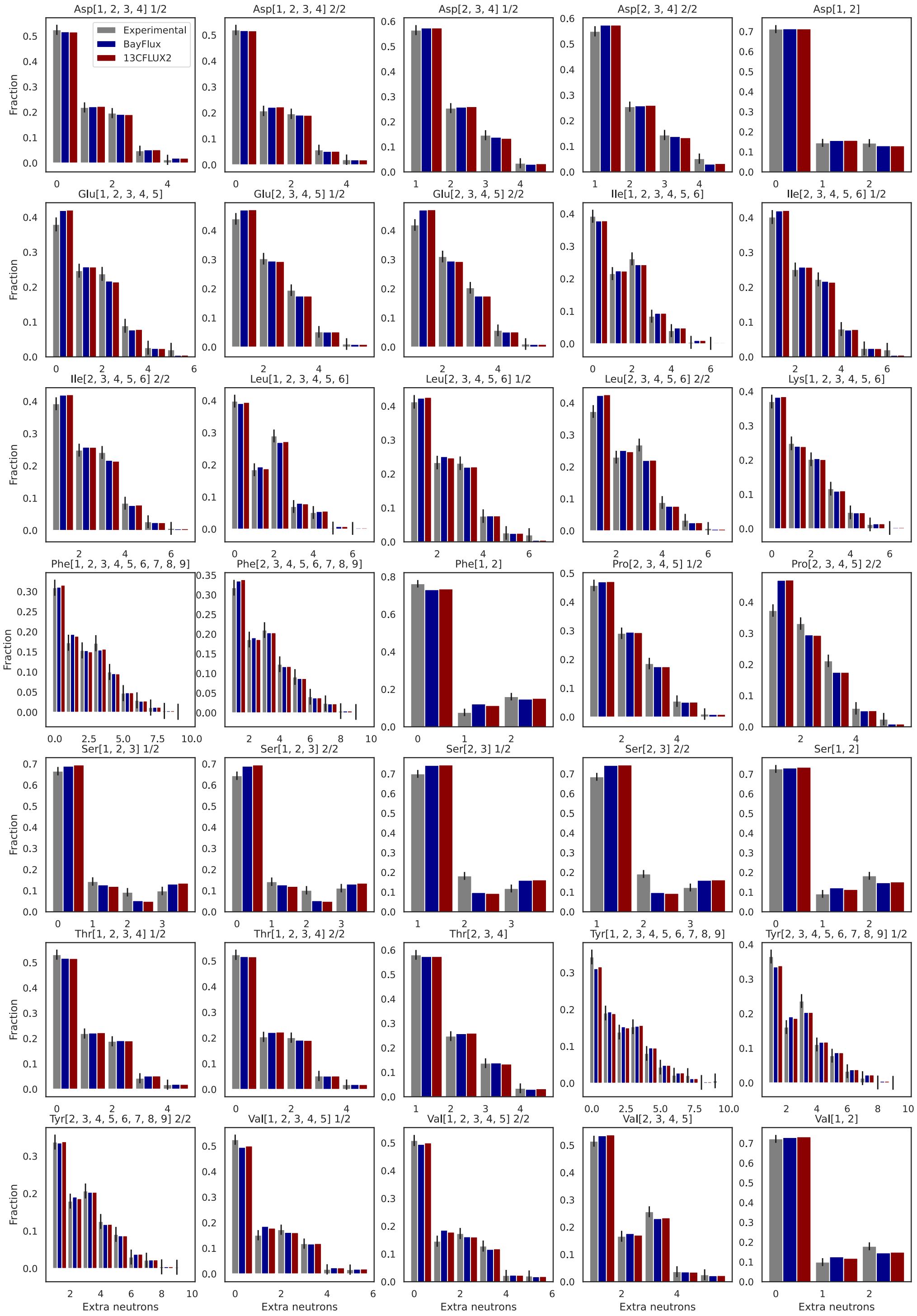


**Fig S1. Fit labeling patterns are the same for both approaches, BayFlux and 13CFLUX2**. Mass distributions vectors (MDVs) for experimental data (grey), the fits for the best sample for BayFlux (dark blue, ten million samples), and the fits for 13CFLUX2 (red) are closer together than the standard deviation of the experimental error. Units for horizontal axis are the number of extra neutrons in the measured metabolite [54].

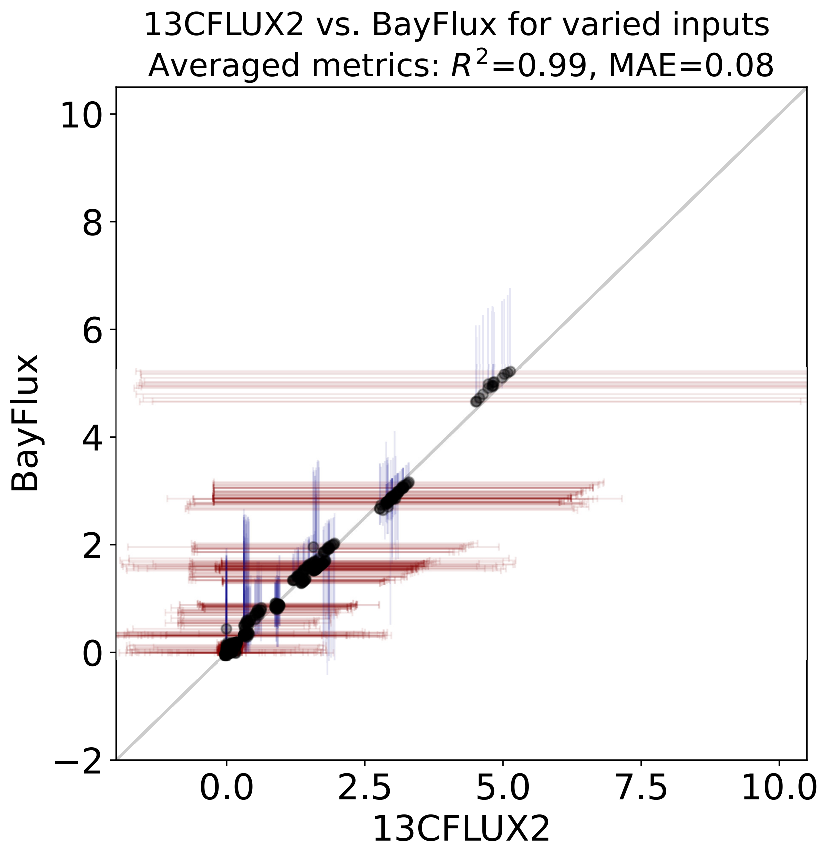


**Fig S2. Flux results for BayFlux and 13CFLUX2 are essentially the same for fifteen different inputs.** The best BayFlux sample (dark blue, ten million samples, y axis) and its credible interval overlap with the 13CFLUX2 best fit (in red, x axis) and its confidence interval for all fifteen different inputs and fluxes. The fifteen different inputs were obtained by randomly changing the exchange fluxes, but keeping the same labeling profiles for the metabolites. Both axes are in units of mM/gDW/h. We set the 13CFLUX2 confidence bounds that could not be determined to the overall maximum BayFlux bounds for comparison purposes.

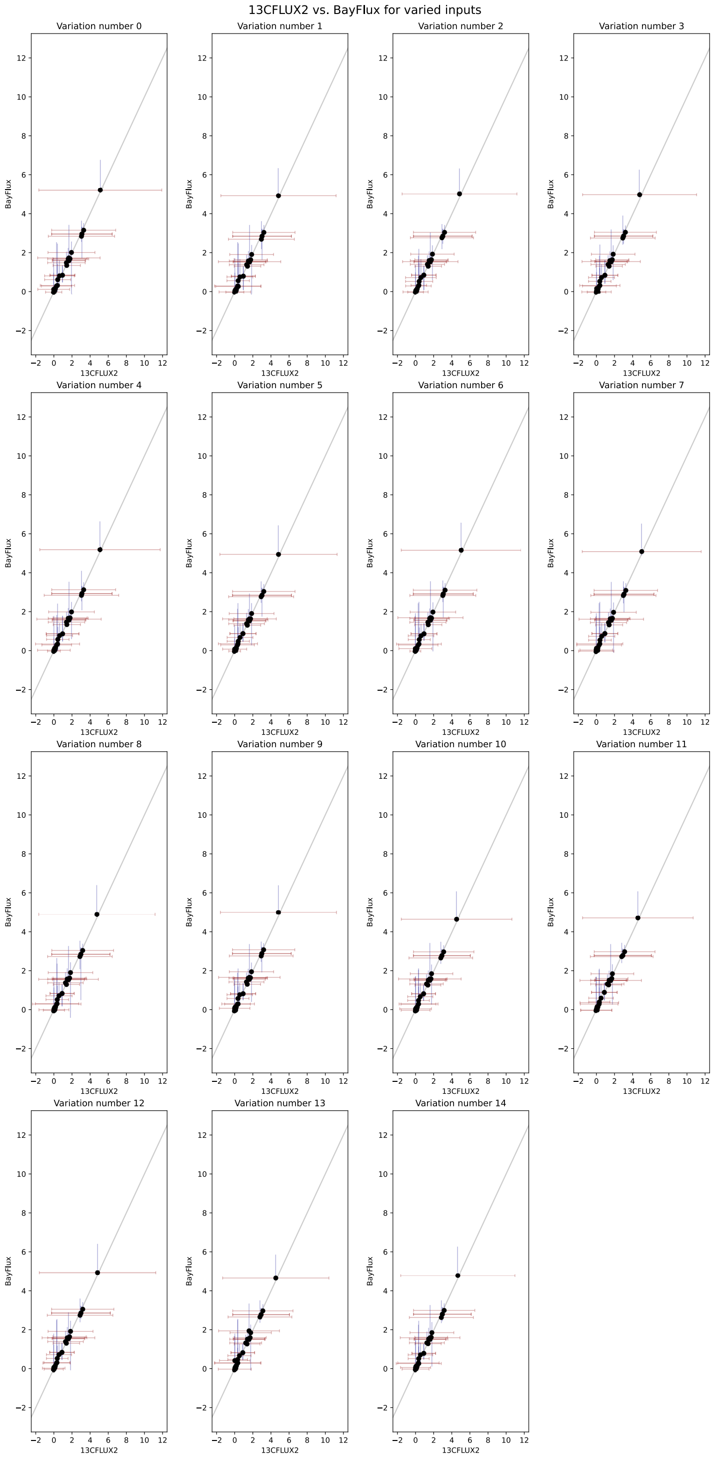


**Fig S3.** 13CFLUX2 vs. BayFlux for 15 different inputs: Best Sample out of 10 million from BayFlux (in dark blue) vs. 13CFLUX2 (in red). We set the 13CFLUX2 confidence bounds that could not be determined to the overall maximum BAYFLUX bounds for comparison purposes.

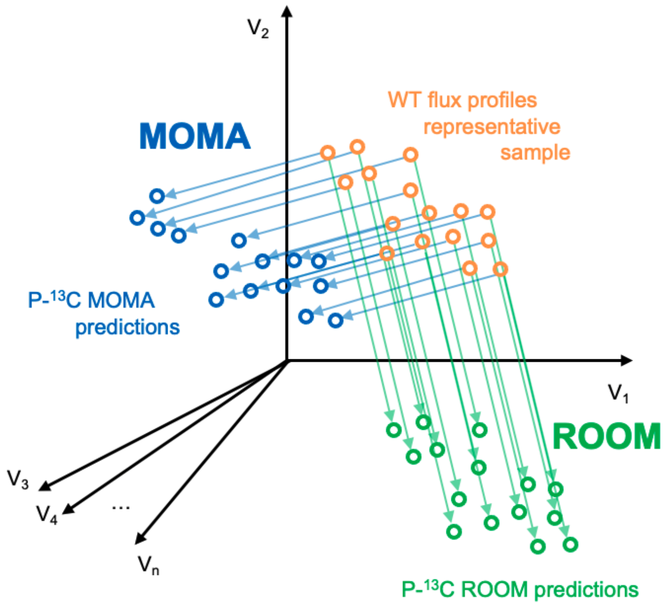


**Fig S4.** P-$^{13}$C MOMA and P-$^{13}$C ROOM work by first computing the WT flux profile distribution using BayFlux (in orange). Each dot corresponds to a flux profile (*i.e.* reaction fluxes for a full genome-scale model) in the flux phase space. A representative set of flux profiles for this base flux profile distribution is obtained through sampling (orange dots). For each of these flux profiles, a new flux profile is computed by using MOMA (in blue) or ROOM (in green) to predict the resulting flux after a knockout. The fact that BayFlux produces a flux profile distribution results in MOMA and ROOM predicting distributions of flux profiles.
